# Invasive dedifferentiated melanoma cells inhibit JAK1-STAT3-driven actomyosin contractility of human fibroblastic reticular cells of the lymph node

**DOI:** 10.1101/2021.03.25.436913

**Authors:** Christopher Rovera, Ilona Berestjuk, Margaux Lecacheur, Serena Diazzi, Aude Mallavialle, Jean Albrengues, Cédric Gaggioli, Christophe Girard, Thierry Passeron, Marcel Deckert, Sophie Tartare-Deckert, Virginie Prod’homme

**Affiliations:** Université Côte d’Azur, Inserm, C3M, Nice, France; Team 11, Equipe labellisée Ligue Contre le Cancer 2016, Nice, France; Université Côte d’Azur, CNRS, Inserm, IRCAN, Nice, France; Université Côte d’Azur, Inserm, C3M, Team 12, Nice, France; Université Côte d’Azur, Department of Dermatology, CHU Nice, Nice, France

**Keywords:** fibroblastic reticular cells, contractility, lymph nodes, melanoma, phenotype plasticity

## Abstract

Fibroblastic reticular cells (FRC) are immunologically specialized fibroblasts controlling the size and microarchitecture of the lymph node (LN), partly through their contractile properties. Swelling is a hallmark of tumor-draining LN in lymphophilic cancers such as cutaneous melanoma, a very aggressive and heterogeneous tumor with high risk of early metastasis. Melanoma cells can dynamically switch between melanocytic proliferative and dedifferentiated mesenchymal-like invasive phenotypes, which are characterized by distinct transcriptional signatures. Melanoma secreted cues, such as extracellular vesicles, growth factors or proinflammatory cytokines, promote LN stroma remodeling and metastatic spreading. But how FRC integrate these pro-metastatic signals and modulate their contractile functions remains poorly characterized. Here, we show that factors secreted by dedifferentiated melanoma cells, but not by melanocytic cells, strongly inhibit FRC actomyosin-dependent contractile forces by decreasing the activity of the RHOA-ROCK pathway and the mechano-responsive transcriptional co-activator YAP, leading to a decrease in F-actin stress fibers and cell elongation. Transcriptional profiling and biochemical analyses indicate that FRC actomyosin cytoskeleton relaxation is driven by inhibition of JAK1 and its downstream transcription factor STAT3, and is associated with increased FRC proliferation and activation. Interestingly, dedifferentiated melanoma cells reduce FRC contractility *in vitro* independently of extracellular vesicle secretion. These data show that FRC are specifically modulated by proteins secreted by invasive dedifferentiated melanoma cells and suggest that melanoma-derived cues could modulate the biomechanical properties of distant LN before metastatic invasion. They also highlight that JAK1-STAT3 and YAP signaling pathways contribute to the maintenance of the spontaneous contractility of resting human FRC.

## Introduction

Melanoma is a very aggressive skin cancer due to its high propensity to metastasis and pronounced intratumor heterogeneity. Four genomic subtypes of cutaneous melanoma have been defined based on the mutational pattern in *BRAF, RAS, NF1*, or none of these (1). During metastatic progression, melanoma cells are highly plastic and dynamically switch between proliferative and invasive phenotypes associated with distinct differentiation states ranging from melanocytic to dedifferentiated (2–5). The proliferative melanocytic state is characterized by high expression of the melanocyte lineage-specific Microphthalmia-Associated Transcription Factor (MITF) and low expression of the tyrosine kinase receptor AXL, whereas the invasive dedifferentiated state shows low expression of MITF and high expression of AXL. Dedifferentiated melanoma cells display a mesenchymal-like phenotype associated with drug resistance (6–8). Whether the melanocytic and dedifferentiated melanoma cell populations differ in their ability to communicate with the metastatic stromal host niche remains poorly understood.

Cutaneous melanoma is a cancer with an inherent potential for lymph node (LN) colonization, an event contributing to systemic metastasis (9–11). Melanoma cells secrete extracellular vesicles (12,13) and soluble factors (14) migrating to the pre-metastatic LN and conditioning immune cells (15), lymphatic endothelial cells (14,16) and fibroblastic reticular cells (FRC) (17). This reprogramming of the LN microenvironment creates a favorable niche that supports metastatic development.

FRC are immunospecialized myofibroblasts of lymphoid organs characterized as CD31^-^Podoplanin^+^ (PDPN) mesenchymal cells. They form a network in close contact with T cells and dendritic cells (DCs) in the LN paracortical area and regulate immune cell recruitment, survival and activation (18). With a notable spontaneous contractibility (19–21) and the secretion and remodeling of a dense reticular network of conduits (18,21), FRC control the LN elasticity and microarchitecture. PDPN was shown to drive the actomyosin contractility of FRC through its binding to proteins of the ezrin, radixin and moesin (ERM) family, leading to RHOA activation (19,20,22,23). During an immune response, migratory DCs expressing high levels of C-type lectin-like receptor 2 (CLEC2) are recruited to the LN. CLEC2 binding to PDPN on FRC dismantles the PDPN-ERM interaction, inhibiting actomyosin contractility and resulting in FRC stretching allowing rapid LN expansion (19,20).

In other tissues, like the skin, quiescent fibroblasts are not spontaneously contractile but can be converted into contractile myofibroblasts by factors secreted during wound healing or tumor progression, such as TGF-β or IL-6 family cytokines (24). Factors secreted by tumor cells transform fibroblasts in the tumor vicinity into cancer associated fibroblasts (CAF) (25). CAF are characterized by a contractile phenotype and the high expression of markers like α-smooth muscle actin (α-SMA, *ACTA2* gene), platelet derived growth factor receptor-α (PDGFR-α) and PDGFR-β, fibroblast activation protein α (FAP), fibroblast-specific protein-1 (FSP1, *S100A4* gene), fibronectin (FN) and a FN isoform containing the EDA domain (EDA-FN), vimentin (VIM) or secreted protein acidic and rich in cysteine (SPARC) (25). In CAF, actomyosin contractility is driven by RHO and RHO-kinase (ROCK) signaling, leading to an increased phosphorylation of the myosin light-chain 2 (MLC2) (26), and by the activation of the mechano-responsive yes1 associated transcriptional regulator (YAP) (27,28). Cytokines from the IL-6 family (IL-6, LIF, OSM) have been shown to induce RHO-ROCK-dependent CAF contractility through the GP130 (IL6ST)-JAK1-STAT3 pathway (29,30). TGF-β increases actomyosin contractility in fibroblasts by promoting LIF expression, which subsequently epigenetically activates JAK-STAT signaling (31). Actomyosin contraction in CAF is associated with cell shape remodeling, increased F-actin stress fibers and drives force-mediated matrix remodeling (26,27).

Malignant LN colonization is preceded by pre-metastatic LN swelling (16,17) but it remains incompletely understood whether (and how) tumor-derived cues alter FRC specific contractile properties during pre-metastatic LN reprogramming. In this study, we characterized the modulation of the actomyosin contractility of isolated human FRC treated with factors secreted by melanocytic or dedifferentiated melanoma cells and investigated the underlying signaling pathways and the nature of the tumoral factor(s) responsible.

## Materials and Methods

### Isolation and culture of primary fibroblasts and CAF

Primary human LN Fibroblasts (LN-F) (#2530, ScienCell) were amplified in Fibroblast Medium (#2301, ScienCell) supplemented with 1% Fibroblast Growth Supplement (#2352, ScienCell), 10% FBS (Gibco) and 100 μg/ml Penicillin/Streptomycin solution (P/S, Gibco). Skin Fibroblasts (Skin-F) were isolated as described previously (32). Melanoma skin or LN clinical specimens (n = 9) were obtained with written informed consent from patients, in accordance with the Declaration of Helsinki, and the study was approved by local ethic committees (Nice Hospital Center and University Côte d’Azur). Briefly, the sample was cut into small pieces and digested with collagenase and dispase. Following filtration of large debris, the solution was serial centrifuged and the final pellet was re-suspended in Dulbecco’s modified Eagle Medium (DMEM) supplemented with 10% FBS and seeded in a tissue culture dish. After 30 min, CAF have adhered to the dish whilst other cellular types in suspension were discarded. The identity of CAF was validated by the expression of classical markers by qRT-PCR and flow cytometry. Skin-F and CAF were cultured in DMEM with 100 μg/ml P/S and 10% FBS. All fibroblasts were starved for 5 to 7 days in DMEM 0.5% FBS (control medium) before any experiment and were then stimulated every two days with control medium or melanoma CM supplemented with 0.5% FBS. They were used until passage 10. In some experiments, cells were treated with 2 ng/ml TGF-β1 (#11343160, ImmunoTools), 2 ng/ml LIF (#300-05, PeproTech) or 10 μM Ruxolitinib (#S1378, Selleckchem), Y-27632 (#S1049, Selleckchem) or SB431542 (#S1067, Selleckchem).

### Melanoma cell culture

Melanoma cell lines were obtained as previously described (6, 33–35). Short-term cultures of patient melanoma cells MM029, MM074 and MM099 were kindly provided by J.-C. Marine and were described elsewhere (8). Patient melanoma cells MNC1 were described previously (35). Cell lines were authentified for mutations by PCR and transcriptional profile by qRT-PCR and Immunoblotting. Cells were cultured in DMEM supplemented with 7% FBS and P/S. All cells were routinely tested for the absence of mycoplasma by PCR.

### Preparation of conditioned media (CM) and mass spectrometry analysis

Melanoma cells were grown until 80% confluent in full growth medium. Then, cells were washed with PBS and cultured for 24h in medium without FBS. The culture supernatant was harvested and filtered (0.45 μM) to eliminate cellular debris. CM prepared from 501Mel and 1205Lu melanoma cell lines were analyzed by mass spectrometry as described (35). In some experiments, EVs and EV-depleted CM were prepared by differential ultracentrifugation as described (35), CM was incubated for 5 min at 95°C, or treated with 100 μg/ml Proteinase K (ThermoFisher) overnight to degrade proteins and then with 1 mM PMSF (Sigma) to inactivate the Proteinase K. In other experiments, CM was filtrated through different membranes with MW cut-offs of 100, 50, 30, 10 and 3 kDa (Amicon Ultra-4, Merck Millipore) to fractionate the proteins according to their size. Resulting CM, or EVs, were aliquoted and frozen at −80°C until used.

### Collagen matrix remodeling assays

Fibroblasts (7.5×10^3^ cells) were embedded in 30 μl of a collagen I (#354249, Corning) - matrigel (#E1270, Sigma) mix and seeded in glass-bottom 96-well plate (#P96G-1.5-5-F, MatTek). Once the gel was set (1h at 37°C), it was overlaid with 100 μl control medium or CM supplemented with 0.5% FBS (with vehicle, cytokines or inhibitors). The matrix was photographed every 1 to 3 days. The respective diameters of the well and matrix were measured using ImageJ software. The percentage of matrix contraction was calculated as followed: 100 - 100 x (matrix area / well area).

### Proliferation assays

Proliferation was measured by a MTS conversion assay using the CellTiter 96 Aqueous Non-Radioactive Cell Proliferation kit (#G5421, Promega) according to the manufacturer’s instructions, or by cell counting.

### RNAi studies

The siRNAs targeting STAT3#1 (#HA13744330, Sigma), STAT3#2 (#HA13744332, Sigma), JAK1 (#SASI_Hs01_00174612, Sigma), JAK2 (#145131, ThermoFisher) and YAP (#HSS115942, ThermoFisher) were used with the respective control siRNAs:

MISSION siRNA Universal Negative Control#1 (Sigma) and a siRNA against the Luciferase Reporter (sense: CGUACGCGGAAUACUUCGATT, antisense: UCGAAGUAUUCCGCGUAC GTT) (ThermoFisher). Transfection of siRNA was carried out using Lipofectamine RNAiMAX (#13778150, ThermoFisher) at a final concentration of 50 nM. Cells were assayed at 2 days post transfection.

### Real-time quantitative PCR

Total RNA was extracted using NucleoSpin RNA Plus kit (#740984.50, Macherey-Nagel). Reverse transcription was performed with the High capacity cDNA Reverse Transcription kit (#4368814, Applied Biosystems). Quantitative PCR was performed using the Platinum SYBR Green qPCR Supermix (#11558656, FisherScientific) with the StepOnePlus System (Applied Biosystems). Relative mRNA levels were determined using the 2ΔΔ*C*t method and *ACTB, GAPDH, HPRT* and *PPIA* as housekeeping genes.

### Microarray experiment

LN-F were cultured for 48h with control medium or 1205Lu CM. RNAs from 4 different experiments (biological quadruplicate) were extracted as described above and analyzed on SurePrint G3 Human Gene Expression 8×60K v2 Microarrays (#G4851B, Agilent Technologies) as previously described (36). Data analyses were performed using R. We used the Bioconductor package array QualityMetrics and custom R scripts for quality control. Additional analyses were done using Bioconductor package Limma. Data were normalized using the quantile method. A linear modeling approach was used to calculate log ratios, moderated *t*-statistics, and *P*-values for each comparison of interest. *P*-values were adjusted for multiple testing using the Benjamini–Hochberg method that controls the false discovery rate.

### Analysis of functional enrichment in gene signatures in microarray data

After raw data processing, the 25,000 most expressed genes in LN-F (average mean expression ≥ 6) were analyzed. To identify pathways down-regulated by 1205Lu CM treatment, we selected the genes down-regulated with LogFC ≤ −0.5 in 1205Lu-reprogrammed LN-F and submitted these genes to Gene Set Enrichment Analysis (GSEA). To analyze genes involved in the regulation of the STAT3 pathway, we identified genes common to the “GO negative regulation of receptor signaling pathway via stat” pathway, and differentially expressed with an absolute LogFC ≥ 0.4. Results were presented in heatmaps generated with Prism 8 (Graphpad) or Phantasus (https://genome.ifmo.ru/phantasus) softwares.

### Immunoblots

Cell lysates were prepared using RIPA buffer (#20-188, Millipore) supplemented with protease and phosphatase inhibitors (#A32959, ThermoFisher). Proteins were separated by SDS-PAGE and were transferred onto PVDF membranes (GE Healthcare) for immunoblot analysis. Membranes were incubated with the primary antibody overnight, washed and then incubated with the peroxidase-conjugated secondary antibody for 1h. The following primary antibodies were used (diluted 1:1,000): α-SMA (#ab5694, Abcam), CD63 (#NBP2-42225, Biotechne), PDPN (#337001, Biolegend), α-Tubulin (#3873), CD9 (#13174), CD81 (#56039), PDGFR-β (#4564), ERM (#3142), P-ERM (T567/T564/T558) (#3149), P-MLC2 (S19) (#3671), STAT3 (#9139), P-STAT3 (Y705) (#9145), YAP (#4912) (Cell Signaling Technology) and FAP (#sc-100528), FN (#sc-9068), HSP60 (#sc-1722), HSP70 (#sc-66048), MLC2 (#sc-15370) (Santa Cruz Biotechnology). Blots were developed with a chemiluminescence system (ECL, #RPN2106, GE Healthcare).

### RHOA assays

RHOA activity was measured on LN-F lysates using the RHOA G-LISA Activation Assay kit (#BK124, Cytoskeleton). Data are the mean ± SEM of arbitrary units of RHOA activity.

### Immunofluorescence analysis

LN-F were grown 4 days on coverslips coated with collagen or on synthetic hydrogels of 2.8 kPa or 0.2 kPa stiffness made as described (6). Then, cells were fixed in 4% paraformaldehyde, permeabilized in PBS 0.3% Triton for 20 min, blocked in PBS 0.1% Triton, 5% Goat Serum for 1h and stained overnight at 4°C with antibodies targeting YAP (#sc-271134, Santa Cruz Biotechnology) and P-MLC2 (S19) (#3671, Cell Signaling Technology) primary antibodies diluted 1:100 in PBS 0.1% Triton, 2% Goat Serum. Following 1h incubation with Alexa Fluor-conjugated secondary antibodies (1:1000, ThermoFisher) and Texas Red- or Alexa Fluor 488-conjugated Phalloidin (1:200, ThermoFisher), nuclei were stained with Hoechst (#H1399, ThermoFisher) and coverslips were mounted in ProLong diamond antifade (#P36961, ThermoFisher). Images were acquired on a wide-field microscope (Leica DM5500B, 40X magnification) or a confocal microscope (Nikon Eclipse Ti, 20X or 40X magnification). Images were analyzed with the ImageJ software to quantify the cell shape index, the mean fluorescence per cell (Integrated density) and the nuclear/cytosolic ratio of YAP, assessed with the fluorescence intensity of nucleus (delimited by Hoechst staining) and cytosol.

### Flow cytometry

Cells were washed in PBS and incubated at 4°C for 30 min in PBS 2% FBS, 2 mM EDTA with CD31-AF488 (#303109), CD35-PE (#333405) and PDPN-APC (#337021) or control isotypes antibodies (Biolegend). Then, cells were washed in PBS and analyzed with BD FACSCANTO II cytometer (BD Biosciences) and the FlowJo software.

### Statistical analysis

Statistical analysis was performed using Prism 8 (GraphPad). Unpaired two-tailed Mann-Whitney tests were used for statistical comparisons between two groups and Kruskal-Wallis tests with Dunn’s post-tests or two-way analysis of variance (ANOVA) tests with Sidak’s post-tests to compare three or more groups. Histogram plots and curves represent mean ± SEM and Violin plots represent median +/- quartiles. P values ≤ 0.05 were considered statistically significant.

### Data availability

The experimental data from microarray have been deposited in the NCBI Gene Expression Omnibus (GEO) database under the series record GSE157355.

## Results

### LN fibroblasts harbor a CAF-like phenotype associated with spontaneous cell contractility

To investigate the interactions between melanoma cells and FRC, we employed primary fibroblasts isolated from human LN. Because several mesenchymal subsets coexist in the LN (21,37), we first characterized these primary LN fibroblasts (LN-F) using microarray profiling, qRT-PCR and flow cytometry (Supplementary Fig. S1). We found that genes expressed by LN-F were typical of FRC (*PECAM1^-^, PDPN^+^, CR2^-^, MADCAM1^-^*), the most abundant LN mesenchymal subset. Flow cytometry analysis further validated the expression of FRC markers (CD31^-^, PDPN^+^, CD35^-^).

FRC exhibit spontaneous contractility (19–21). When embedded in a collagen-based three-dimensional matrix, LN-F isolated from 4 different donors demonstrated basal force-mediated matrix remodeling, driven by the ROCK-actomyosin pathway and inhibited by the ROCK inhibitor Y-27632 (**Fig. 1A**). LN-F displayed the same high propensity to drive collagen matrix remodeling as primary CAF isolated from skin or LN melanomas (**Fig. 1B**). However, primary skin fibroblasts (skin-F) were not able to contract the collagen matrix unless activated with TGF-β to a CAF-like phenotype. In contrast, LN-F spontaneous contraction was not modulated by TGF-β. As for LN-F, contractile properties of TGF-β-activated skin-F and CAF were dependent of the ROCK-actomyosin pathway and inhibited by Y-27632. In agreement with the collagen matrix remodeling results, LN-F contractile activity was associated with higher basal levels of F-actin stress fibers and YAP nuclear localization compared to skin-F (**Fig. 1C** and **D**). However, in contrast to skin-F, F-actin fibers and YAP nuclear localization were not increased by TGF-β treatment in LN-F. Analysis by qRT-PCR revealed that several CAF markers like *ACTA2* (α-SMA), *FAP*, *EDA-FN* or *SPARC* were highly expressed in LN-F compared to skin-F, suggesting that resting LN-F share many properties with CAF (**Fig. 1E**). Immunoblot analysis also showed TGF-β-independent high expression of FN, PDGFR-β, FAP and α-SMA in resting LN-F, to levels equivalent or higher than TGF-β-activated skin-F (**Fig. 1F**). This is consistent with the notion that high α-SMA expression is associated with fibroblast contractility (38). Collectively, these results validate the FRC signature of LN-F and unveil that quiescent LN-F display some phenotypic and functional properties of CAF. Our aim was then to investigate whether, and how, melanoma cells modulate fibroblast contractile properties of the LN metastatic niche.

**Figure 1.**
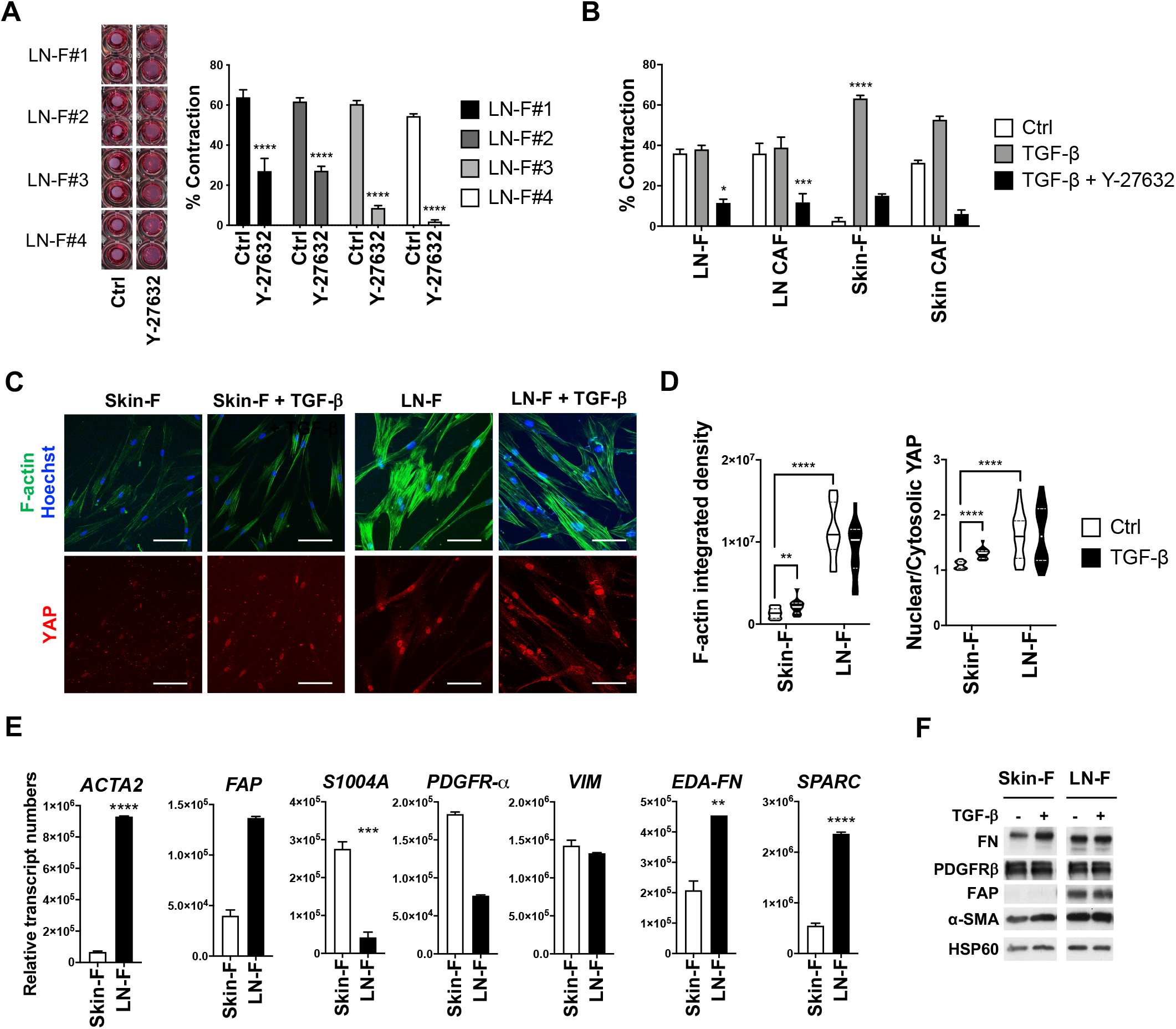
LN-F display a CAF-like phenotype associated with spontaneous contraction and expression of CAF markers. (A) Pictures of 3D collagen matrices containing LN-F from 4 donors (#1 to #4) treated with control medium (Ctrl) or the ROCK inhibitor Y-27632 (10 μM) for 8 days (left). Quantification of the LN-F-mediated collagen matrix contraction (in quadruplicate; mean +/- SEM; p-Val (****)<0.0001)) (right). (B) Matrix contraction induced by LN-F, skin-F or CAFs isolated from skin or LN melanomas and treated with control medium (Ctrl), TGF-β (2 ng/ml) or Y-27632 (10 μM) for 7 days (n = 2 to 7 for each cell type, in triplicate; mean +/- SEM; p-Val (*)<0.05, (***)<0.001, (****)<0.0001). (C) Fluorescence pictures showing F-actin fibers, cell nuclei (Hoechst) and YAP localization in Skin-F and LN-F grown for 4 days in control medium or with TGF-β (2 ng/ml) (Scale bar = 100 μm). (D) Quantification of F-actin integrated density (n = 19 cells; median +/- quartiles) and YAP localization (n = 22 cells; median +/- quartiles); p-Val (**)<0.01, (****)<0.0001). (E) Quantification by qRT-PCR of the expression of CAF markers in LN-F and skin-F (n = 2, in duplicate; mean +/- SEM; p-Val (**)<0.01, (***)<0.001, (****)<0.0001). (F) Immunoblotting of CAF markers (FN, PDGFRβ, FAP and α-SMA) in Skin-F and LN-F cultured for 4 days in control medium (Ctrl) or treated by TGF-β (2 ng/ml). HSP60 is used as a loading control.

### Factors secreted by invasive dedifferentiated melanoma cells with a MITF^low^ AXL^high^ signature inhibit LN-F contractility

Melanoma cells harbor transcriptional states ranging from melanocytic (MITF^high^ AXL^low^) to dedifferentiated (MITF^low^ AXL^high^) (2–5). To compare the ability of these melanoma sub-populations to modulate LN-F contractility, we selected several melanocytic (501Mel, MeWo, SK-MEL-28, WM164) and dedifferentiated (1205Lu, WM793, WM2032, SBcl2) cell lines identified by qRT-PCR analysis of MITF and AXL gene expression (Supplementary Fig. S2A). Then, to model melanoma distant reprogramming of fibroblasts in the pre-metastatic LN, we treated LN-F with conditioned medium (CM) harvested from melanoma cell cultures and containing the factors secreted by melanoma cells. Factors secreted by tumor cells, like TGF-β, are known to induce fibroblast actomyosin contraction, as during CAF differentiation (25). Strikingly, we observed that CM secreted by dedifferentiated melanoma cell lines (MITF^low^ AXL^high^) drastically inhibited LN-F spontaneous contractility, while melanocytic melanoma cell lines (MITF^high^ AXL^low^) showed no effect (**Fig. 2A**). Similar results were obtained with short-term patient melanoma cultures as CM prepared from dedifferentiated MNC1, MM029 and MM099 cells inhibited LN-F contractility. In contrast, CM from the melanocytic MM074 cells displayed no statistically inhibitory effect (**Fig. 2B** and supplementary S2B). Together, our results demonstrate that the ability to modulate LN-F contractility is strongly correlated to the dedifferentiated MITF^low^ signature of invasive melanoma cells (**Fig. 2C**). Next we analyzed whether inhibition of contractility requires prolonged LN-F exposure to tumoral factors. After 8 days treatment, LN-F in collagen matrix settings recovered their spontaneous contractility upon 1205Lu CM withdrawal (**Fig. 2D**). Thus, melanoma cell-induced inhibition of LN-F contraction is a reversible process and requires the presence of factors secreted by dedifferentiated melanoma cell to be maintained.

**Figure 2.**
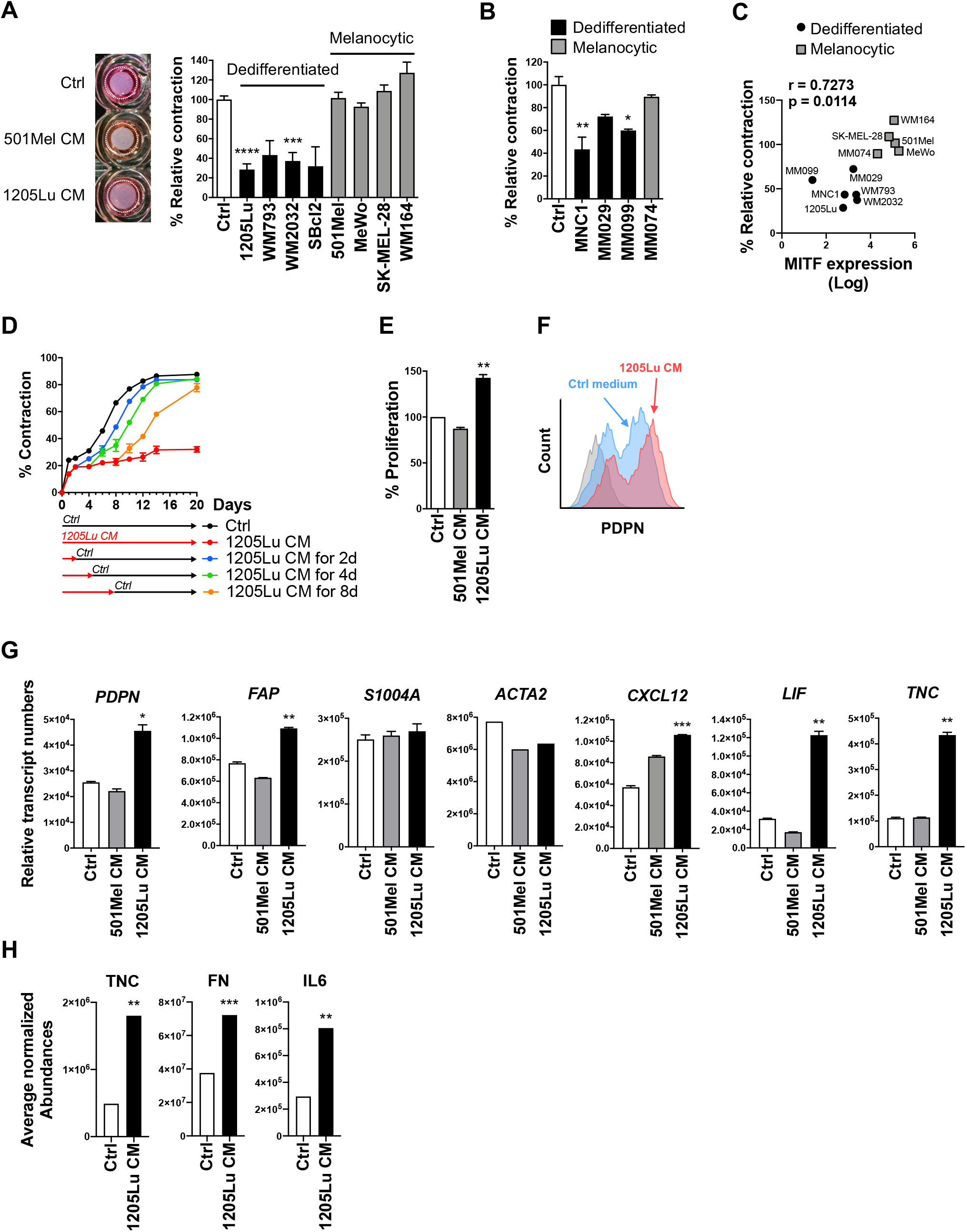
Factors secreted by MITF^low^ AXL^high^ dedifferentiated melanoma cells inhibit LN-F contractility and induce LN-F proliferation and activation. (A) Representative picture (left) and quantification (right) of LN-F-mediated contraction of a collagen matrix after 7 days culture with control (Ctrl) medium or CM from dedifferentiated MITF^low^ AXL^high^ or melanocytic MITF^high^ AXL^low^ melanoma cell lines (n = 3, in triplicate; mean +/- SEM; p-Val (***)<0.001, (****)<0.0001). (B) LN-F-mediated collagen matrix contraction after 7 days incubation with CM from dedifferentiated MITF^low^ AXL^high^ or melanocytic MITF^high^ AXL^low^ short term patient melanoma cells (n = 3, in triplicate; mean +/- SEM; p-Val (*)<0.05, (**)<0.01). (C) Plot showing MITF gene expression by melanoma cells (Log) versus the % of LN-F-mediated collagen matrix contraction shown in (A) and (B) (Spearman correlation analysis). (D) Time-lapse analysis of LN-F-mediated collagen matrix contraction (n = 2, in quadruplicate; mean +/- SEM). The 1205Lu CM is replaced by Ctrl medium after 2, 4 or 8 days. (E) Proliferation of LN-F incubated 6 days with ctrl medium, 501Mel CM or 1205Lu CM (n = 2, in triplicate; mean +/- SEM; p-Val (**)<0.01). (F) Representative flow cytometry analysis of PDPN surface expression on LN-F cultured for 5 days with 1205Lu CM (in red) or ctrl medium (in blue). Staining with a control isotype mAb is shown in grey (n = 4). (G) qRT-PCR analysis of fibroblast activation markers expressed by LN-F cultured for 2 days with ctrl medium, 501Mel CM or 1205Lu CM (n = 2, in duplicate; Mean +/- SEM; p-Val (*)<0.05, (**)<0.01, (***)<0.001). (H) Detection by mass spectrometry of TNC, FN and IL6 secreted by LN-F treated for 7 days with ctrl medium or 1205Lu CM (n = 2, in triplicate; Mean +/- SEM; p-Val (**)<0.01, (***)<0.001).

To get a more comprehensive view of the effect of factors secreted by melanocytic or dedifferentiated melanomas on LN-F reprogramming, we carried out the study with two representative melanoma cell lines displaying the melanocytic MITF^high^ AXL^low^ phenotype (501Mel) or the dedifferentiated MITF^low^ AXL^high^ phenotype (1205Lu). During an immune challenge, FRC activation is associated with proliferation, up-regulation of PDPN and other markers of fibroblast activation, and disruption of the RHOA-ROCK-mediated actomyosin cytoskeleton contraction due to CLEC2 interaction with PDPN (19,20,39). Similarly, 1205Lu-derived CM (1205Lu CM) stimulated LN-F proliferation (**Fig. 2E**) and increased the cell surface expression of PDPN observed by flow cytometry whereas 501Mel CM had no effect (**Fig. 2F**). qRT-PCR analysis showed that expression of several markers of fibroblast activation like *PDPN*, *FAP*, *CXCL12*, *LIF* and *TNC* was also up-regulated by 1205Lu CM while *S100A4* (FSP1) and *ACTA2* (α-SMA) levels were not affected by treatment with 501Mel or 1205Lu CM (**Fig. 2G**). Mass spectrometry (MS) analysis also revealed that 1205Lu-activated LN-F secreted more extracellular matrix proteins (TNC, FN) and cytokines (IL-6), validating the functional activation of LN-F (**Fig. 2H**). These results indicate that the inhibition of FRC contractility induced by dedifferentiated melanoma CM is a functional response of activated LN-F, similar to the FRC behavior observed during immunization. Importantly, our observations suggest that factors secreted by invasive dedifferentiated and proliferative melanocytic melanoma cells do not share the same abilities to reprogram LN-F.

The propensity of fibroblasts to contract a collagen matrix is regulated by forces generated by the actomyosin cytoskeleton (26,27). To learn more about the modulation of LN-F contractility by dedifferentiated melanoma CM, we examined by immunofluorescence the F-actin organization and phosphorylation of the ROCK substrate MLC2 in LN-F treated or not with 501Mel CM or 1205Lu CM. The ROCK inhibitor Y-27632 was used as a control (**Fig. 3A** and **B**). We observed similar changes related to inhibition of actomyosin contraction (27) in LN-F treated with Y-27632 or 1205Lu CM, like decreased MLC2 S19 phosphorylation, less F-actin fibers and the remodeling of the cell morphology from a stellate to a fusiform shape.

**Figure 3.**
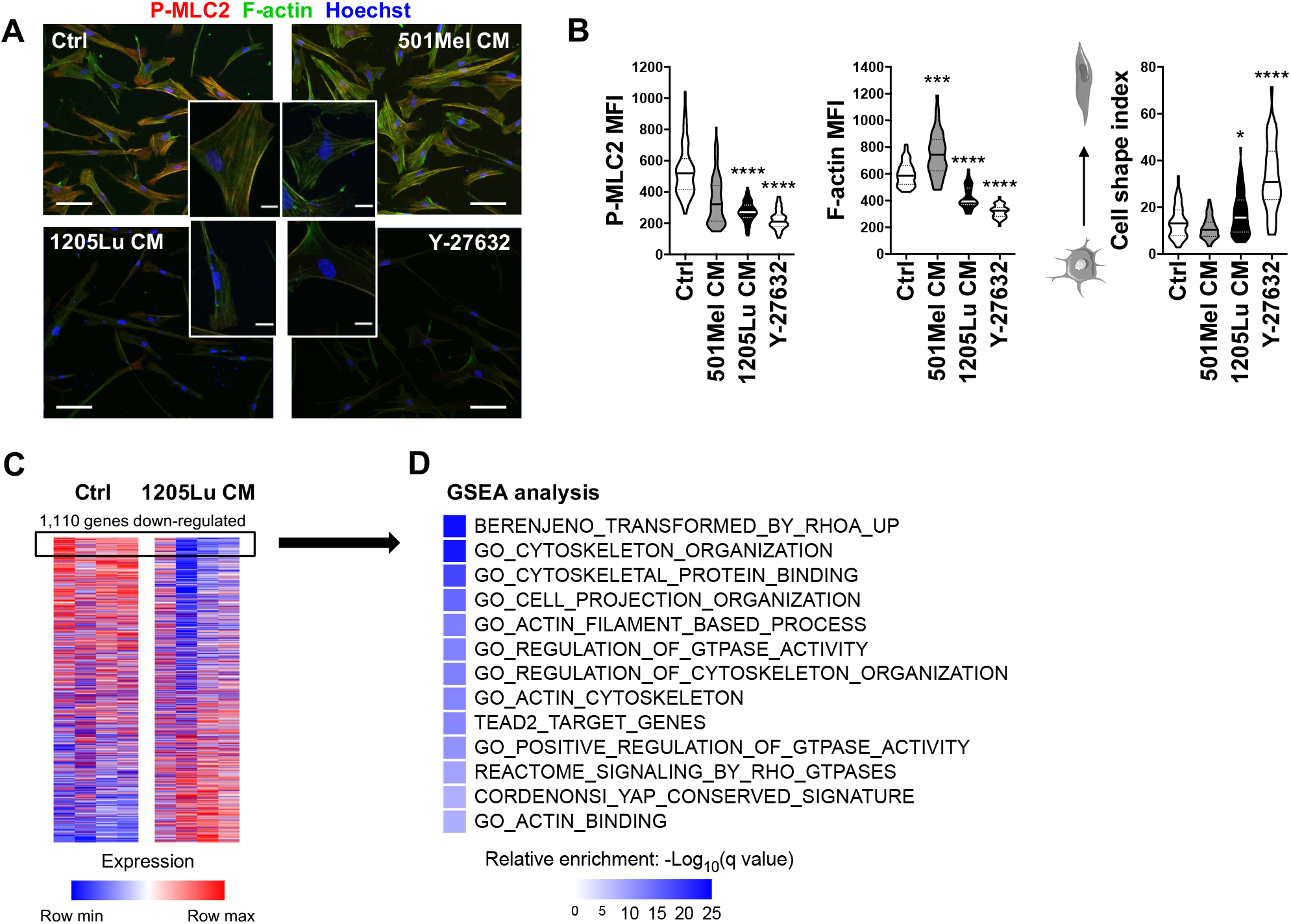
Factors secreted by the MITF^low^ AXL^high^ dedifferentiated 1205Lu cell line inhibit the LN-F actomyosin cytoskeleton contraction. (A) Fluorescence staining of F-actin fibers, MLC2 phosphorylation (P-MLC2) and cell nuclei (Hoechst) in LN-F grown for 7 days in control (Ctrl) medium or with CM from the dedifferentiated 1205Lu cell line or the melanocytic 501Mel cell line (Scale bar = 100 μm). A zoom on a single cell is shown for each condition (Scale bar = 20 μm). (B) Quantification of mean fluorescence intensities (MFI) of active pS19 MLC2 (P-MLC2), F-actin fibers and the cell shape index (n = 45 cells; median +/- quartiles; p-Val (*)<0.05, (***)<0.001, (****)<0.0001). (C) Microarray-based gene expression profiling in Ctrl and 1205Lu CM-treated LN-F. The 1,110 most down-regulated genes in 1205Lu-reprogrammed LN-F (with LogFC ≤ - 0.5) are surrounded and submitted to Gene Set Enrichment Analysis (GSEA) in (D).

In order to identify molecular pathways responsible for inhibition of LN-F contractility by dedifferentiated melanoma CM, we performed microarray-based gene expression profiling on LN-F cultured in control conditions or exposed to 1205Lu CM (**Fig. 3C**). From the 25,000 most expressed LN-F genes, we selected differentially expressed genes (DEGs) with an absolute LogFC ≥ 0.5. We identified 1,113 DEGs up-regulated in 1205Lu CM-treated LN-F and 1,110 DEGs down-regulated, validating LN-F transcriptional reprogramming by the CM from dedifferentiated melanoma cells. Gene Set Enrichment Analysis (GSEA) of the 1,110 down-regulated genes validated the inhibition of pathways related to actin cytoskeleton polymerization, RHOA GTPase activity and pointed towards the regulation of YAP and its co-transcription factor TEAD2 in 1205Lu-reprogrammed LN-F (**Fig. 3D** and Supplementary Fig. S3).

### 1205Lu CM-mediated suppression of LN-F contractility is associated with impaired YAP activity

To validate the inhibition of the YAP pathway identified by GSEA (**Fig. 3D** and **4A**), we investigated the effect of melanocytic and dedifferentiated melanoma CMs on YAP nuclear localization and activation. YAP is sensitive to the cell microenvironment stiffness (28). The stiffness in the LN paracortical area was measured to 0.2 kPa by nanoindentation, and was similar to the one of collagen matrices used to assess LN-F contractility (data not shown). However, in 2D cell culture conditions YAP activity is reduced in such soft hydrogel (28). Thus, we decided to examine the effect of melanoma cell CMs on stiffer hydrogels (2.8 kPa), allowing the optimal visualization of changes in YAP localization and activity.

**Figure 4.**
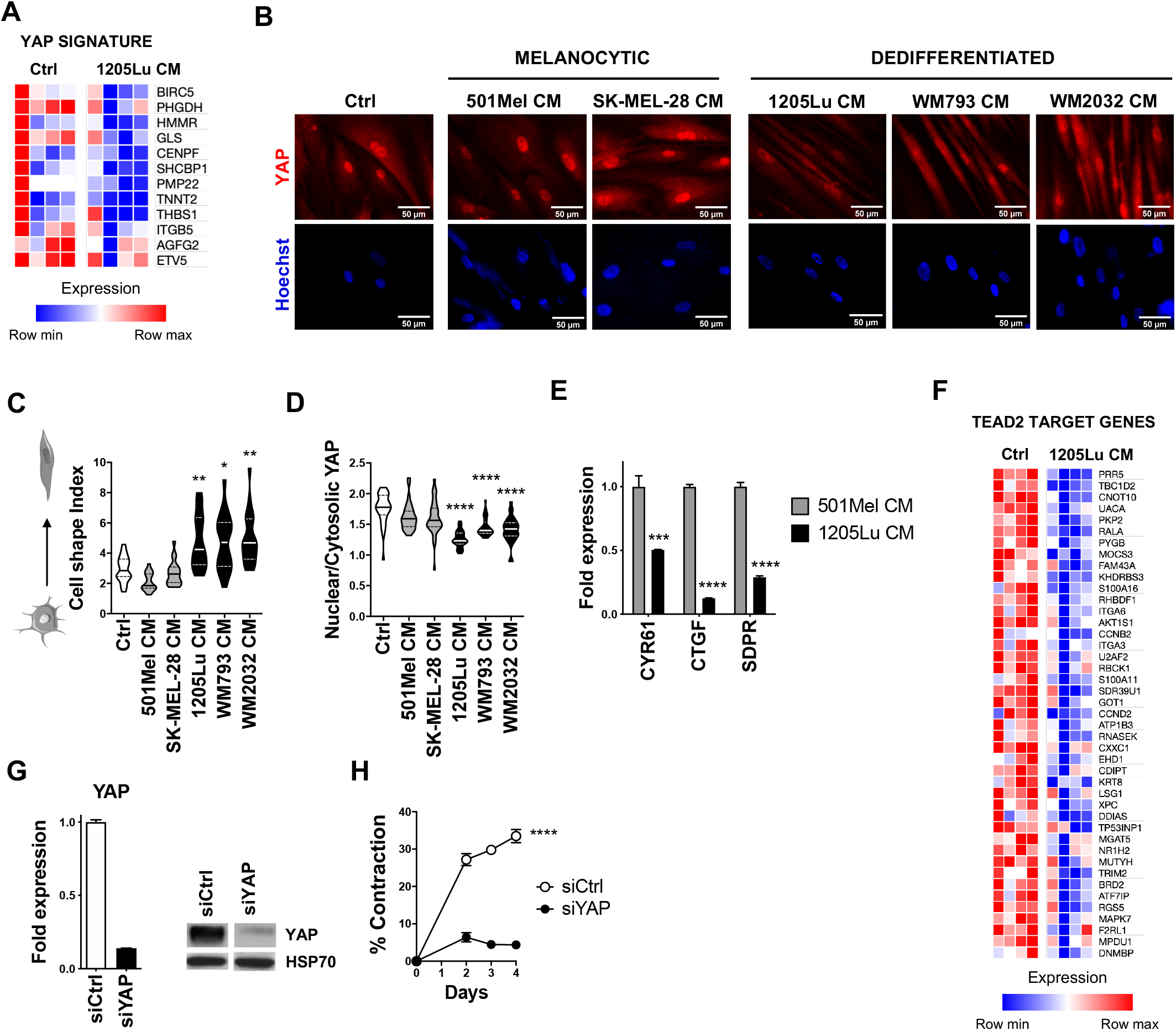
The inhibition of LN-F contractility induced by invasive melanoma CM is associated with decreased YAP activity. (A) Top genes down-regulated in 1205Lu CM-treated LN-F in the Cordenonsi YAP conserved Signature (LogFC ≤ - 0.5). (B) Immunofluorescence analysis of YAP and Hoechst localization in LN-F plated on 2.8 kPa hydrogels and treated 4 days with control (Ctrl) medium, proliferative or invasive melanoma CM (Scale bar = 50 μm). (C) Analysis of the cell shape index of LN-F plated on 2.8 kPa hydrogels and treated 4 days with ctrl medium, proliferative or invasive melanoma CM (n = 20 cells; median +/- quartiles; p-Val (*)<0.05, (**)<0.01). (D) Quantification of YAP nuclear and cytosolic localization in LN-F plated on 2.8 kPa hydrogels and treated 4 days with ctrl medium, proliferative or invasive melanoma CM (n = 32 cells; median +/- quartiles; p-Val (****)<0.0001). (E) Quantification by qRT-PCR of the expression of YAP target genes in LN-F plated on 2.8 kPa hydrogels and treated 4 days with 501Mel CM or 1205Lu CM (n = 2, in duplicate; mean +/- SEM; p-Val (***)<0.001, (****)<0.0001) (F) Top TEAD2 target genes down-regulated in 1205Lu CM-treated LN-F (LogFC ≤ - 0.5, pVal ≤ 0.05). (G) Quantification by qRT-PCR (left) and immunoblot (right) of the extinction of YAP expression by siRNA (siYAP) in the experiment shown in F. A ctrl siRNA (siCtrl) is used as a ctrl of transfection (n = 2, in duplicate; mean +/- SEM). (H) Quantification of matrix contraction by LN-F transfected with siCtrl or siYAP (n = 2, in quadruplicate; p-Val (****)<0.0001).

In agreement with the inhibition of LN-F contractility induced by dedifferentiated melanoma cues, immunofluorescence analysis revealed that LN-F incubated with CM from the dedifferentiated 1205Lu, WM793 or WM2032 cells displayed an elongated shape, with more cytosolic YAP compared to control LN-F (**Fig. 4B**-**D**). On the contrary, CM prepared from melanocytic 501Mel or SK-MEL-28 cells induced no significant change on YAP nuclear localization and cell elongation. Consistently, qRT-PCR analysis revealed decreased levels of YAP target genes *CYR61, CTGF* and *SDPR* induced by 1205Lu CM (**Fig. 4E**). We also noticed a decreased expression of genes regulated by the YAP co-transcription factor TEAD2 in LN-F treated with 1205Lu CM (**Fig. 4F**). Although less pronounced, results were similar in LN-F treated with 1205Lu CM on 0.2 kPa hydrogels (Supplementary Fig. S4A-C).

To address the contribution of YAP in LN-F contractility, YAP expression was silenced using a siRNA approach (**Fig. 4G**). The spontaneous LN-F-mediated collagen matrix remodeling was suppressed by YAP knockdown, revealing that YAP was not only a marker of cell contraction, but actually actively controlled LN-F contractility (**Fig. 4H**). Note that YAP depletion by siRNA had no effect on LN-F proliferation (Supplementary Fig. S5A). Together, our data suggest that relaxation of the FRC actomyosin network induced by factors secreted by dedifferentiated melanoma cells is associated with inhibition of YAP-TEAD2 transcriptional activity.

### The JAK1-STAT3 pathway is inhibited by invasive dedifferentiated melanoma CM and is required for LN-F contraction

Next, we sought to determine signaling pathways linking actomyosin cytoskeleton relaxation and YAP inhibition induced by melanoma secreted factors in LN-F. Previous studies have demonstrated that PDPN drives actomyosin contraction of FRC through binding to proteins from the ERM family, which link membrane proteins to the cytoskeleton (22,23). Binding of PDPN to its only known ligand CLEC2 dismantled the PDPN-ERM interaction and inhibited ERM phosphorylation and PDPN-driven actomyosin contraction (19,20). Our first series of data excluded the possible contribution of the CLEC2-PDPN-ERM pathway in melanoma-induced LN-F relaxation. Indeed, expression of CLEC2 was not detected in 1205Lu cells (data not shown) and immunoblot analysis showed that ERM phosphorylation was not affected in 1205Lu CM-treated LN-F (**Fig. 5A**). However, 1205Lu CM increased PDPN expression, as already shown by flow cytometry in Fig. 2F, and reduced α-SMA protein levels. This is in agreement with LN-F activation and the concomitant inhibition of F-actin stress fiber formation observed in presence of 1205Lu CM (**Fig. 3A** and **B**).

**Figure 5.**
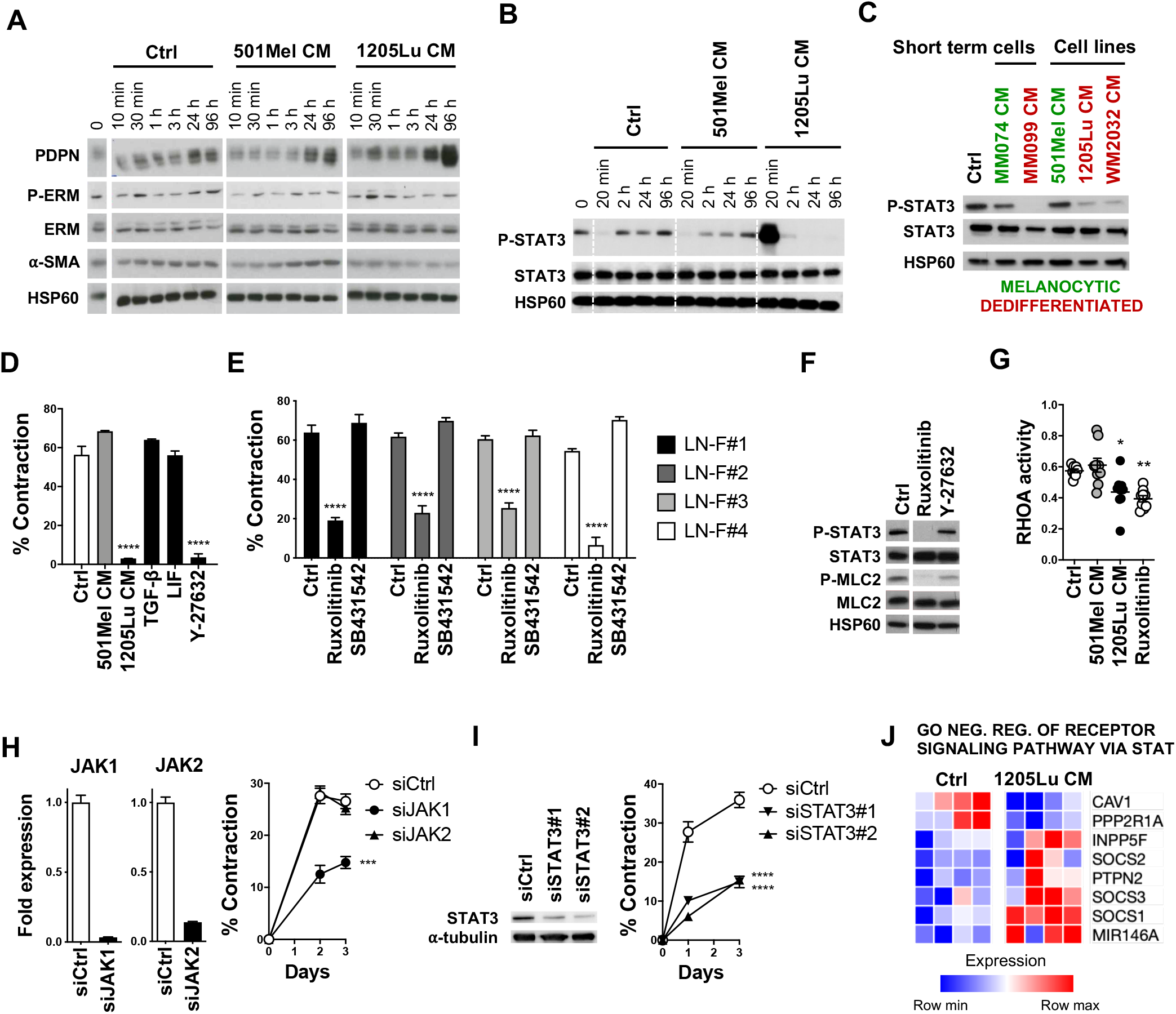
The JAK1-STAT3 pathway is inhibited by invasive melanoma CM and controls LN-F contraction. (A) Immunoblotting of PDPN, P-ERM, ERM and α-SMA in LN-F cultured in control (Ctrl) medium, 501Mel CM or 1205Lu CM for 10 min, 30 min, 1 h, 3 h, 24 h and 96 h. HSP60 is used as a loading control. (B) Immunoblotting of P-STAT3 and STAT3 in LN-F cultured in Ctrl medium, 501Mel CM or 1205Lu CM for 20 min, 2 h, 24 h and 96 h. HSP60 is used as a loading control. (C) Immunoblotting of P-STAT3 and STAT3 in LN-F cultured for 96 h in Ctrl medium or CM from short term isolated melanoma cells (MM074 and MM099) or melanoma cell lines (501Mel, 1205Lu, WM2032). HSP60 is used as a loading control. (D) Matrix contraction by LN-F treated by Ctrl medium, 501Mel CM, 1205Lu CM, TGF-ß1 (2 ng/ml), LIF (2 ng/ml) or Y-27632 (10 μM) for 6 days (n = 3, in triplicate; mean +/- SEM; p-Val (****)<0.0001). (E) Matrix contraction by LN-F treated by 10 μM of TGF-ßR1 inhibitor (SB431542) or Ruxolitinib for 8 days (n = 3, in quadruplicate; mean +/- SEM; p-Val (****)<0.0001). (F) Immunoblotting of P-STAT3, STAT3, P-MLC2 and MLC2 in LN-F treated 4 days by Ruxolitinib (10μM) or Y-27632 (10μM). HSP60 is used as a loading control. (G) RHOA activity measured by G-LISA assay in cell lysates of LN-F treated 4 days with 501Mel CM, 1205Lu CM or Ruxolitinib (10μM) (n = 3, in triplicate; mean +/- SEM; p-Val (*)<0.05, (**)<0.01). (H) qRT-PCR quantification of JAK1 and JAK2 silencing by siRNAs (siJAK1 and siJAK2) (n = 2, in duplicate) (left) and quantification of matrix contraction by LN-F transfected with siJAK1 or siJAK2 (n = 2, in triplicate; mean +/- SEM; p-Val (**)<0.01) (right). A ctrl siRNA (siCtrl) is used as a ctrl of transfection. (I) Immunoblotting of STAT3 and Tubulin-α 2 days after the extinction of STAT3 expression by 2 siRNAs (siSTAT3#1 and siSTAT3#2) (left). Quantification of LN-F-mediated matrix contraction after siRNA silencing of STAT3 (n = 2, in quadruplicate; mean +/- SEM; p-Val (****)<0.0001) (right). siCtrl is used as a control of siRNA transfection. (J) Differentially expressed genes between Ctrl and 1205Lu CM-treated LN-F identified in the GO negative regulation of receptor signaling pathway via STAT.

Because TGF-β and IL-6 family cytokines are known to regulate ROCK-mediated actomyosin contraction in myofibroblasts and CAF (25,29,30), we reasoned that the TGF-β Receptor 1 (TGF-βR1)-SMAD pathway or the GP130-JAK-STAT3 pathway could be involved in LN-F contractility. We previously identified by MS analysis that TGF-β, IL-6 and LIF were highly produced by 1205Lu cells compared to 501Mel cells (35). Indeed, STAT3 phosphorylation was strongly increased after 20 min exposure to 1205Lu CM, but decreased after few hours and remained suppressed the following days (**Fig. 5B**). This indicates that the JAK-STAT3 pathway was specifically and consistently inhibited by factors secreted by dedifferentiated melanoma cells. On the contrary, compared with basal STAT3 phosphorylation observed in LN-F before the addition of fresh medium (time 0), STAT3 phosphorylation was decreased 20 min after the addition of control (Ctrl) medium or 501Mel CM, but returned to basal levels in 2 h. These results suggest that resting LN-F activated an autocrine JAK-STAT3 loop, which was interrupted when the LN-F supernatant was replaced by fresh medium. Indeed, we identified by MS that resting LN-F secrete basal levels of IL6 and LIF (**Fig. 2H** and data not shown). Sustained inhibition of STAT3 phosphorylation was also observed after 4 days treatment with CM prepared from other dedifferentiated melanoma cells, such as the WM2032 cell line or the short-term MM099 cells. In contrast, after treatment with CM from the melanocytic short-term MM074 melanoma cells, levels of STAT3 phosphorylation in LN-F were comparable to the control (**Fig. 5C**).

We next investigated the consequences of the modulation of TGF-βR1-SMAD and GP130-JAK-STAT3 pathways on LN-F spontaneous contractility. Treatment with TGF-β or LIF had no effect on LN-F basal contractility (**Fig. 5D**), nor the inhibition of TGF-βR1 signaling by SB431542 (**Fig. 5E**). However, the JAK1/2 inhibitor Ruxolitinib strongly inhibited the spontaneous contraction of LN-F isolated from 4 different donors (**Fig. 5E**). Note that treatment with Ruxolitinib, SB431542 or Y-27632 had no effect on LN-F proliferation (Supplementary Fig. S5B). Importantly, immunoblot analysis revealed that Ruxolitinib inhibited both STAT3 and MLC2 phosphorylation, linking the JAK-STAT pathway to actomyosin contraction in LN-F (**Fig. 5F**). Indeed, similar to the effect of 1205Lu CM, Ruxolitinib inhibited RHOA activity in LN-F, suggesting that suppression of actomyosin contraction induced by 1205Lu CM was mediated through RHOA inhibition by the JAK-STAT pathway (**Fig. 5G**). As Ruxolitinib inhibits both JAK1 and JAK2 kinase activities, we used siRNAs against JAK1 (siJAK1) or JAK2 (siJAK2) to discriminate which JAK kinase(s) was involved in LN-F contraction. Because JAK1 depletion, but not JAK2, inhibited LN-F-mediated collagen matrix contraction, we identified JAK1 as the main JAK involved in LN-F contraction (**Fig. 5H**). Those results were further confirmed with the observation that STAT3 extinction by two different siRNA inhibited LN-F collagen matrix contraction, suggesting that JAK1 controls LN-F contractility through STAT3 (**Fig. 5I**). JAK1 or STAT3 depletion by siRNAs had no effect on LN-F proliferation (Supplementary Fig. S5A). Looking back to the GSEA analysis of genes down-regulated in 1205Lu CM-treated LN-F compared to Ctrl (**Fig. 3D**), it seemed surprising that the JAK-STAT pathway modulation could not be identified. So we repeated the GSEA analysis with the most down-regulated, but also up-regulated genes from the microarray (**Fig. 5J**). Indeed, several genes strongly up-regulated in 1205Lu CM-treated LN-F were related to the negative regulation of the JAK-STAT pathway, such as the MIR146A microRNA (40), members of the SOCS (Suppressor Of Cytokine Signaling) family (41) and the phosphatases PTPN2 (Protein Tyrosine Phosphatase Non-Receptor Type 2) (42) and INPP5F (Inositol Polyphosphate-5-Phosphatase F) (43). Together, our results reveal the contribution of JAK1-STAT3 signaling pathway in 1205Lu-mediated suppression of LN-F contraction.

### The JAK1-STAT3 pathway controls YAP activity and the actin cytoskeleton polymerization

We next questioned whether JAK1-STAT3 inhibition impacts on YAP function and ROCK-MLC2-actomyosin cytoskeleton remodeling in LN-F. Fibroblasts were treated with the JAK1/2 inhibitor Ruxolitinib or the ROCK inhibitor Y-27632 and analyzed by fluorescence microscopy (**Fig. 6A**). In agreement with the role of the RHOA-ROCK pathway in YAP activity and actomyosin polymerization (28), we found that pharmacological inhibition of ROCK induced YAP cytosolic relocation (**Fig. 6B**), decreased F-actin stress fibers (**Fig. 6C**) and LN-F stretching (**Fig. 6D**). Similar observations were made when cells were treated with Ruxolitinib, suggesting that JAK1 is critical to maintain nuclear YAP and actomyosin network tension. Consistently, JAK1 silencing by siRNA (siJAK1) induced YAP cytosolic relocation and inhibited the formation of actin filaments (**Fig. 6E-G**). Interestingly, similar modifications were observed after STAT3 extinction (siSTAT3), suggesting that JAK1 effects on YAP and actomyosin are mediated by STAT3. Accordingly, expression of the YAP target genes *CTGF* and *CYR61* was inhibited after STAT3 knockdown by siRNA (**Fig. 6H**).

**Figure 6.**
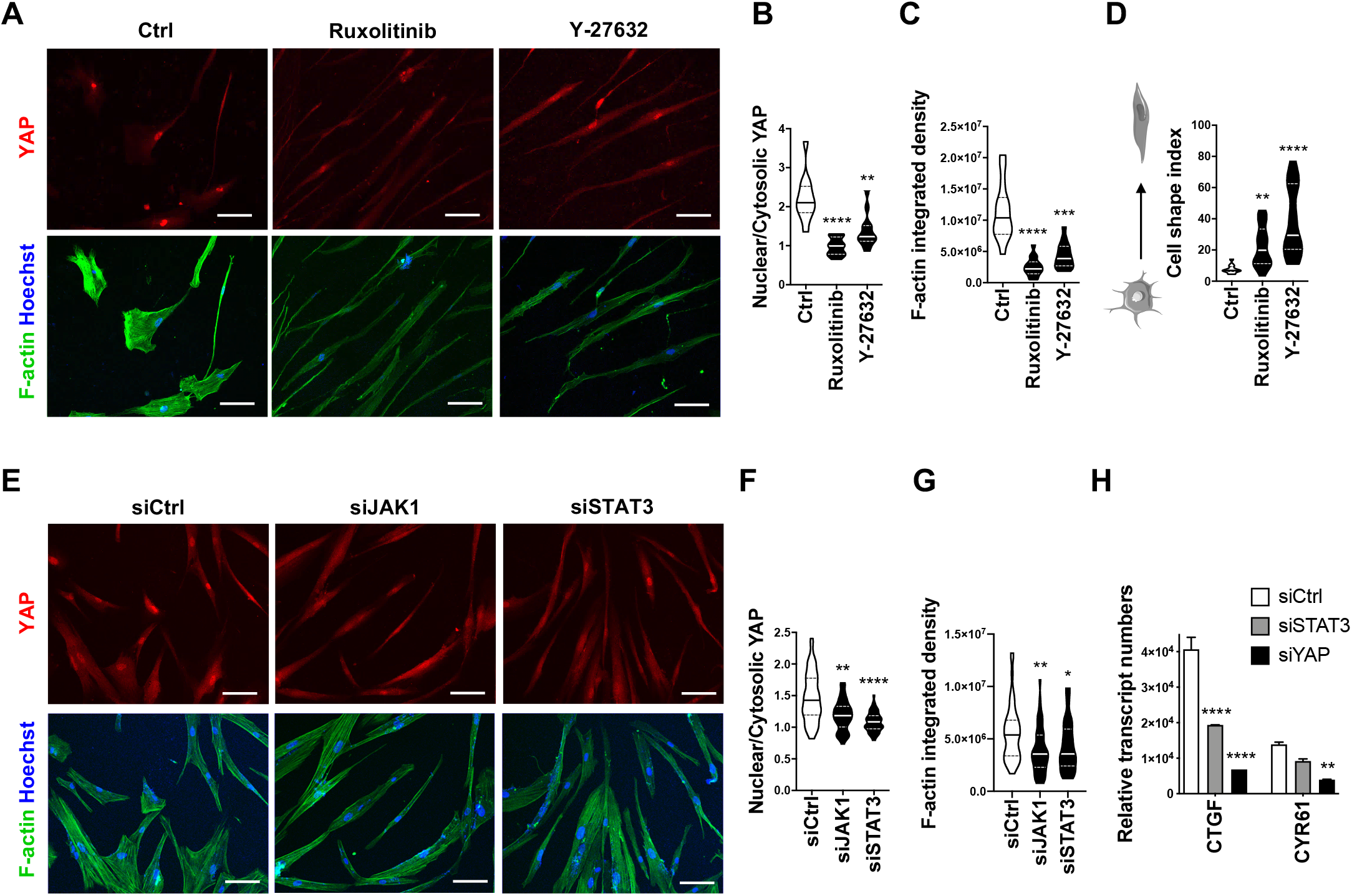
The JAK1-STAT3 pathway controls YAP and polymerization of the actin cytoskeleton. (A) Immunofluorescence analysis of YAP, F-actin fibers and Hoechst in LN-F treated for 4 days with Ruxolitinib (10 μM) or Y-27632 (10 μM) (Scale bar = 100 μM). (B) Quantification of YAP localization (n = 13 cells; median +/- quartiles; p-Val (**)<0.01, (****)<0.0001). (C) Quantification of F-actin integrated density (n = 15 cells; median +/- quartiles; p-Val (***)<0.001, (****)<0.0001). (D) Quantification of the cell shape index (n = 20 cells; median +/- quartiles; p-Val (**)<0.01, (****)<0.0001). (E) Immunofluorescence analysis of YAP, F-actin fibers and Hoechst in LN-F depleted from YAP (siYAP) or STAT3 (siSTAT3) expression by siRNA. The siCtrl siRNA is used as a control of transfection (Scale bar = 100 μM). (F) Quantification of YAP localization (n = 50 cells; median +/- quartiles; p-Val (**)<0.01, (****)<0.0001). (G) Quantification of F-actin integrated density (n = 44 cells; median +/- quartiles; p-Val (*)<0.05, (**)<0.01). (H) qRT-PCR analysis of the expression of YAP target genes (n = 2, in duplicate; mean +/- SEM; p-Val (**)<0.01, (****)<0.0001).

Therefore, after deciphering the signaling pathway linking factors secreted by dedifferentiated melanoma cells to the relaxation of the LN-F actomyosin cytoskeleton, we focused our attention on the nature of these tumoral cues.

### Invasive dedifferentiated melanoma cells release soluble proteins that inhibit LN-F contractility

The CM contains all the factors secreted by melanoma cells, including soluble proteins and lipids and extracellular vesicles (EVs) such as exosomes. EVs are known to promote LN niche formation and distal metastatic tumor growth (12,13). Knowing that 1205Lu cells actually secreted more EVs than 501Mel cells (data not shown), we first investigated if EVs had any effect on LN-F contractility. The 1205Lu CM was depleted from EVs by ultracentrifugation and analyzed by immunoblot to control the enrichment of EV-specific CD9, CD63 and CD81 tetraspanins and the chaperone protein HSP70 in the EV fraction and their absence in the EV-depleted 1205Lu CM fraction (**Fig. 7A**). The LN-F-mediated collagen matrix contraction was inhibited to similar levels after incubation with the EV-depleted 1205Lu CM or the complete 1205Lu CM, while incubation with concentrated 1205Lu EVs (5 μg/ml) did not modulate LN-F spontaneous contractility (**Fig. 7B**). These results excluding any involvement of melanoma EVs in the regulation of LN-F contractility, we turned our attention towards proteins and lipids secreted independently from EVs. Proteins are more sensitive to heat than lipids. As the 1205Lu CM heated to 95°C for 5 min partly lost its ability to inhibit LN-F cell contraction, we therefore hypothesized that proteins, rather than lipids, were responsible for the 1205Lu CM-inhibition of LN-F contractility (**Fig. 7C**). To validate this idea, we treated the 1205Lu CM by the broad-spectrum serine protease proteinase K (PK) to degrade proteins and then inactivated PK with the serine protease inhibitor PMSF (PhenylMethylSulfonyl Fluoride) before adding the 1205Lu CM on LN-F. The PK-treated 1205Lu CM was no longer able to decrease LN-F-mediated collagen matrix contraction, further confirming the contribution of soluble melanoma-derived proteins in suppression of LN-F contractility (**Fig. 7D**). Finally, to identify more precisely the set of candidate proteins mediating this effect, the 1205Lu CM was filtrated through membranes with distinct molecular weight (MW) cut-offs ranging from 3 kDa to 100 kDa to fractionate the proteins according to their size (**Fig. 7E**). 1205Lu-secreted proteins selected by cut-offs bellow 30 kDa and above 100 kDa did not modify LN-F contractility while proteins selected by cut-offs above 30 kDa retained the ability to inhibit LN-F contraction (**Fig. 7F**). We thus restricted 1205Lu-secreted factors able to regulate LN-F actomyosin polymerization to EV-free proteins with a MW ranging from 30 kDa to 100 kDa.

**Figure 7.**
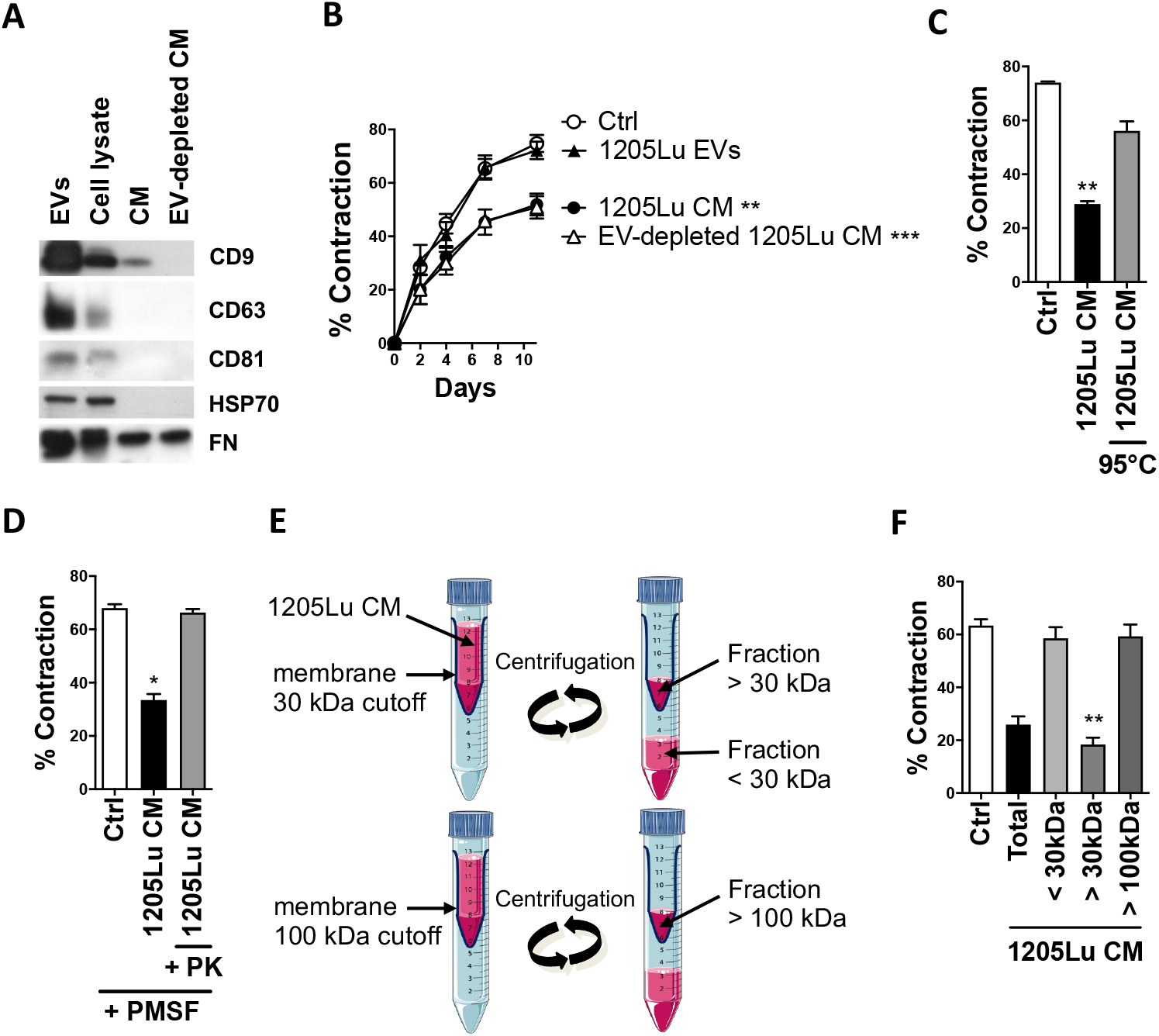
Soluble proteins secreted by dedifferentiated melanoma cells inhibit LN-F contractility. (A) Immunoblot analysis of EV-specific markers CD9, CD63, CD81 and HSP70 in extravesicles (EVs) isolated from the 1205Lu CM (15 μg), in the 1205Lu cell lysate (15 μg), and in the EV-depleted 1205Lu CM. FN is used as a loading control. (B) Time-lapse matrix contraction by LN-F incubated with ctrl medium or samples immunoblotted in A, i.e. 1205Lu CM, EV-depleted 1205Lu CM, and 1205Lu EVs (5 μg/ml) (n = 2, in quadruplicate; mean +/- SEM; p-Val (**)<0.01, (***)<0.001). (C) Matrix contraction by LN-F treated with 1205Lu CM or 95°C heated 1205Lu CM (n = 3, in quadruplicate; mean +/- SEM; p-Val (**)<0.01). (D) Matrix contraction by LN-F treated with Ctrl medium, 1205Lu CM or 1205Lu CM treated with Proteinase K (PK). The proteases were inactivated in the three conditions with phenylmethylsulfonyl fluoride (PMSF) before incubation with LN-F (n = 2, in triplicate; mean +/- SEM; p-Val (*)<0.05). (E) The 1205Lu CM was fractionated by centrifugation using 30 kDa or 100 kDa cut-off membranes. The 1205Lu CM fractions with 1205Lu factors bellow 30 kDa (< 30 kDa), above 30 kDa (> 30 kDa) and above 100 kDa (> 100 kDa) were harvested and tested in the gel contraction assay shown in F. (F) Matrix contraction by LN-F treated with the full 1205Lu CM or 1205Lu fractions bellow 30 kDa (< 30 kDa), above 30 kDa (> 30 kDa) and above 100 kDa (> 100 kDa) (n = 4, in quadruplicate; mean +/- SEM; p-Val (**)<0.01).

## Discussion

Our study provides the first evidence that human FRC are contractile cells similar to murine FRC (19–21). Quiescent FRC display a myofibroblast-like phenotype along with high expression of PDPN, FAP and α-SMA and thus share many properties with CAF. These CAF hallmarks suggest that FRC could play a tumor supportive role in the LN metastatic microenvironment. Indeed, PDPN was identified as a CAF marker in a variety of malignancies and is associated with metastasis and poor prognosis (44,45). PDPN expression on CAF favors force-mediated matrix remodeling through the activation of the RHO-ROCK pathway and promotes cancer cell invasion (45). In a recent study in breast cancer patients, metastatic LN were shown to be enriched in a CAF subpopulation inducing cancer cell invasion and exhibiting similar markers (PDPN^high^, FAP^high^, α-SMA^high^, PDGFR-β^high^) to FRC (46), suggesting that LN CAF mostly originated from resident FRC.

Focusing on the signaling pathway(s) driving FRC actomyosin contractility, we provide evidence that the spontaneous contractility of quiescent FRC relies on a basal level of YAP and JAK1-STAT3 activation, and not only on the PDPN-ERM pathway as previously identified in murine FRC. YAP activation was known to reflect the actomyosin contractile state of FRC (20) and CAF (27), and to regulate FRC differentiation during LN development (47). We show here that YAP is not only a marker but also controls actomyosin contractility of differentiated human FRC. YAP and the RHO-ROCK pathway are intimately linked, regulating each other and controlling the actomyosin cytoskeleton plasticity (27,28). We also disclose that inhibition of JAK1 or STAT3 reduces FRC contractility and leads to inhibition of RHOA and YAP activities. Previous studies have shown that JAK1-STAT3 signaling increases ROCK-mediated actomyosin contractility in CAF, and that ROCK signaling induces STAT3 phosphorylation and transcriptional responses. Thus, JAK1-STAT3 and RHO-ROCK are interdependent and cross-regulate each other (30). However, it is still unclear if JAK1-STAT3 controls both YAP and RHOA pathways or if it controls only YAP or RHOA, which then regulate other(s) pathway(s). Collectively, our results support the notion that the RHO-ROCK-driven contractility of human FRC is not only controlled by the PDPN-ERM pathway, but also by signaling pathways shared with CAF.

Tumor cells secrete numerous growth factors and inflammatory factors, such as TGF-β or the IL-6 family cytokines that trigger fibroblast activation and CAF transformation, both processes associated with increased actomyosin contractility. Strikingly, we discover that melanoma cells with a dedifferentiated signature (MITF^low^ AXL^high^) secrete factors drastically suppressing force-driven matrix remodeling by FRC. Factors secreted by dedifferentiated melanoma cells but not by melanocytic melanoma cells, inhibit JAK1-STAT3 signaling, which decreases RHO-ROCK-MLC2 signaling and YAP activity. These pathways are interconnected and may be regulated through feedback mechanisms. The JAK-STAT pathway is a major signaling mechanism for a wide array of cytokine and growth factor receptors. Further investigations are now required to identify the receptor(s) involved in actomyosin relaxation of FRC treated with factors secreted by dedifferentiated melanoma cells. Because CAF activation by tumoral cues is associated with the acquisition of contractile properties, the inhibition of FRC contractility by melanoma CM was unexpected. However, it is part of the FRC activation process, like during an immune challenge (19,20). Indeed we show that FRC stimulated with CM from dedifferentiated melanoma cells are in an activated state. They proliferate more, up-regulate some markers of activation (FAP, PDPN), and increase the secretion of extracellular matrix proteins (FN, TNC) and cytokines (IL-6). Changes in cell contractile potential alter fibroblast proliferation, and inflammation is known to trigger FRC proliferation and PDPN expression, while the specific underlying mechanisms remain unknown (19,20,39). Thus, the increased proliferation that we observe in presence of invasive melanoma secreted factors might have stemmed from dysregulated tension or be independently due to factors secreted by invasive melanoma cells. Interestingly, both BRAF-mutated (1205Lu, WM793, MM099) and NRAS-mutated (WM2032, SBcl2) melanoma dedifferentiated cells were able to inhibit FRC contractility. Melanoma cell plasticity is thought to be a key driver of melanoma progression (2–5). The phenotypic switch to a dedifferentiated MITF^low^ signature is associated with a gain in invasive and metastatic abilities. Our findings further suggest that the dedifferentiated state endows melanoma cells with reprogramming of the LN fibroblastic stroma prior metastatic dissemination. The question remains whether tumor-derived factors from other cancers that spread to the LN, such as breast cancer, have the ability to exert a similar task on FRC contractility and activation.

Culture of FRC with factors secreted by dedifferentiated melanoma cells strikingly phenocopies the prevention of PDPN signaling by the provision of CLEC2, antibody blockade or genetic deficiency (19,20). It induces similar morphological changes with an elongated cell shape, decreased stress fibers, less YAP nuclear activation, but also increased proliferation and PDPN expression. So, while we have excluded the modulation of ERM phosphorylation by melanoma CM, the implication of PDPN in the inhibition of FRC contractility mediated by melanoma CM is still an open question. PDPN can activate the FRC actomyosin machinery independently of its binding to ERM proteins, by engaging a neighboring transmembrane protein (20). Our hypothesis is that this unknown transmembrane protein may drive JAK1 signaling and that binding to its ligand secreted by dedifferentiated melanoma cells could dissociate such interaction. An alternative could be that JAK1 may directly bind to the PDPN cytoplasmic tail. Indeed, JAK1 contains a FERM domain in its N-terminal portion, like ERM proteins, and this domain is responsible for direct binding to the cytoplasmic tail of integral membrane proteins (48). Indeed, PDPN controls a wide range of physiological effects, such as contractility, migration, proliferation or differentiation, probably through distinct molecular partners in diverse cell types (45).

It is established that factors secreted by melanoma cells, including TGF-β, growth factors, or proinflammatory molecules promote tumorigenesis (49) and LN metastasis (14). Melanoma EVs have been particularly described as potent inducers of LN pre-metastatic niches (12,13). However, 1205Lu-EVs did not modulate FRC contractility in our experimental setting and we have restricted candidate factors to EV-free proteins between 30 kDa and 100 kDa. We previously identified by MS that dedifferentiated 1205Lu cells secrete higher levels of cytokines, chemokines, growth factors and matricellular proteins than melanocytic 501Mel cells (34,35,50). However, factors known to affect fibroblast contractility and secreted more abundantly by 1205Lu cells (i.e. TGF-β, IL-6, LIF) did not modulate FRC contractility. In the light of the melanoma cell transcriptional signatures identified (2–5), a detailed analysis of the factors secreted by dedifferentiated and melanocytic subpopulations of melanoma cells is needed to better apprehend their interactions with the components of the tumor microenvironment and identify soluble proteins controlling FRC contractility, proliferation and activation.

FRC contractile phenotype regulates LN swelling and tunes LN immunity (19,20). Factors secreted by cutaneous melanoma may therefore account for the swelling observed in melanoma draining LNs (16,17). One study has analyzed *in vitro* and *in vivo* stromal reprogramming induced by melanoma cells in tumor draining LN using the murine B16.F10 cell line (17). This study reported an increased RHO signaling in murine FRC isolated from B16.F10-draining LN, and in contrast to our work, the enhanced capacity of FRC treated with B16.F10 CM to contract collagen matrices. At present, the reason behind this discrepancy remains unclear. One explanation would be that the murine B16.F10 melanoma cell line, which harbors neither BRaf nor NRas oncogenic mutation, is known to express high levels of MITF despite its metastatic potential, and may control FRC functions differently than MITF^low^ human melanoma cells. To this regard it would be of great importance to compare in a mouse model how melanocytic MITF^low^ and dedifferentiated MITF^high^ melanoma cells reprogram FRC in the LN metastatic niche. Together this underlines that the development of the metastatic disease is a very complex process fueled by intratumor heterogeneity and the mutational context.

LNs from wild-type mice are significantly less stiff and more deformable after immunization or following antibody blockade of PDPN (20), demonstrating that FRC contractility affects LN stiffness. Microenvironment clues such as tissue stiffness might control the back and forth switches of melanoma cells between proliferative melanocytic and invasive dedifferentiated states to drive disease progression. We can hypothesize that a softer pre-metastatic LN microenvironment could then favor the switch of invasive dedifferentiated melanoma cells reaching the LN to proliferative melanocytic cells able to form new metastatic foci. The increased elasticity of the reticular FRC network conferred by actomyosin relaxation could also allow more LN stretching to give space for metastatic development without LN breakdown. Loss of FRC contractility also leads to significant changes in the homeostasis and spacing of FRC and T cells, with profound consequences for the LN microarchitecture and the population expansion of antigen-specific T cells following immunization. FRC control immune cell survival and proliferation through cell contacts and secretion of immunoregulatory molecules (18). Spacing of FRC and metastatic cells could also regulate tumoral development with similar mechanisms. Reprogramming the biomechanical and immune properties of the LN through loss of FRC contractility thus may be essential for the survival and development of metastatic cells in the LN niche.

In conclusion, we have identified a new mechanism by which invasive melanoma cells can reprogram the function of FRC from the LN niche and deciphered the underlying intracellular signaling pathways involved in FRC cytoskeleton relaxation. Our work illustrates that FRC activation and actomyosin relaxation in LN might be a prognostic marker of the melanoma invasive capacities, suggesting that the microarchitecture of the FRC reticular network and PDPN expression could be examined in tumor-draining LN at diagnosis. Blocking the distant communication between invasive melanoma cells and FRC may inhibit the formation of a permissive biomechanical and immunological LN niche, thereby impinging on lymphatic metastasis.

## Acknowledgements

We thank J.-C. Marine for short-term cultures of melanoma cells, and the “Microscopie Imagerie Côte d’Azur” (MICA) GIS-IBISA platform supported by the GIS IBiSA, Conseil Départemental 06 and Région Provence Alpes Côte d’Azur. We acknowledge M. Gesson and M. Irondelle from the C3M imaging facility for their helpful advice.

## References

1. Cancer Genome Atlas N. Genomic Classification of Cutaneous Melanoma. Cell 2015;161(7):1681–96 doi 10.1016/j.cell.2015.05.044.

2. Arozarena I, Wellbrock C. Phenotype plasticity as enabler of melanoma progression and therapy resistance. Nat Rev Cancer 2019;19(7):377–91 doi 10.1038/s41568-019-0154-4.

3. Hoek KS, Eichhoff OM, Schlegel NC, Dobbeling U, Kobert N, Schaerer L, et al. In vivo switching of human melanoma cells between proliferative and invasive states. Cancer Res 2008;68(3):650–6 doi 10.1158/0008-5472.CAN-07-2491.

4. Rambow F, Rogiers A, Marin-Bejar O, Aibar S, Femel J, Dewaele M, et al. Toward Minimal Residual Disease-Directed Therapy in Melanoma. Cell 2018;174(4):843–55 e19 doi 10.1016/j.cell.2018.06.025.

5. Wouters J, Kalender-Atak Z, Minnoye L, Spanier KI, De Waegeneer M, Bravo Gonzalez-Blas C, et al. Robust gene expression programs underlie recurrent cell states and phenotype switching in melanoma. Nat Cell Biol 2020;22(8):986–98 doi 10.1038/s41556-020-0547-3.

6. Girard CA, Lecacheur M, Ben Jouira R, Berestjuk I, Diazzi S, Prod’homme V, et al. A Feed-Forward Mechanosignaling Loop Confers Resistance to Therapies Targeting the MAPK Pathway in BRAF-Mutant Melanoma. Cancer Res 2020;80(10):1927–41 doi 10.1158/0008-5472.CAN-19-2914.

7. Shaffer SM, Dunagin MC, Torborg SR, Torre EA, Emert B, Krepler C, et al. Rare cell variability and drug-induced reprogramming as a mode of cancer drug resistance. Nature 2017;546(7658):431–5 doi 10.1038/nature22794.

8. Verfaillie A, Imrichova H, Atak ZK, Dewaele M, Rambow F, Hulselmans G, et al. Decoding the regulatory landscape of melanoma reveals TEADS as regulators of the invasive cell state. Nat Commun 2015;6:6683 doi 10.1038/ncomms7683.

9. Brown M, Assen FP, Leithner A, Abe J, Schachner H, Asfour G, et al. Lymph node blood vessels provide exit routes for metastatic tumor cell dissemination in mice. Science 2018;359(6382):1408–11 doi 10.1126/science.aal3662.

10. Pereira ER, Kedrin D, Seano G, Gautier O, Meijer EFJ, Jones D, et al. Lymph node metastases can invade local blood vessels, exit the node, and colonize distant organs in mice. Science 2018;359(6382):1403–7 doi 10.1126/science.aal3622.

11. Ubellacker JM, Tasdogan A, Ramesh V, Shen B, Mitchell EC, Martin-Sandoval MS, et al. Lymph protects metastasizing melanoma cells from ferroptosis. Nature 2020;585(7823):113–8 doi 10.1038/s41586-020-2623-z.

12. Hood JL, San RS, Wickline SA. Exosomes released by melanoma cells prepare sentinel lymph nodes for tumor metastasis. Cancer Res 2011; 71(11):3792–801 doi 10.1158/0008-5472.CAN-10-4455.

13. Peinado H, Aleckovic M, Lavotshkin S, Matei I, Costa-Silva B, Moreno-Bueno G, et al. Melanoma exosomes educate bone marrow progenitor cells toward a pro-metastatic phenotype through MET. Nat Med 2012;18(6):883–91 doi 10.1038/nm.2753.

14. Olmeda D, Cerezo-Wallis D, Riveiro-Falkenbach E, Pennacchi PC, Contreras-Alcalde M, Ibarz N, et al. Whole-body imaging of lymphovascular niches identifies pre-metastatic roles of midkine. Nature 2017;546(7660):676–80 doi 10.1038/nature22977.

15. Cochran AJ, Huang RR, Lee J, Itakura E, Leong SP, Essner R. Tumour-induced immune modulation of sentinel lymph nodes. Nat Rev Immunol 2006;6(9):659–70 doi 10.1038/nri1919.

16. Commerford CD, Dieterich LC, He Y, Hell T, Montoya-Zegarra JA, Noerrelykke SF, et al. Mechanisms of Tumor-Induced Lymphovascular Niche Formation in Draining Lymph Nodes. Cell Rep 2018;25(13):3554–63 e4 doi 10.1016/j.celrep.2018.12.002.

17. Riedel A, Shorthouse D, Haas L, Hall BA, Shields J. Tumor-induced stromal reprogramming drives lymph node transformation. Nat Immunol 2016;17(9):1118–27 doi 10.1038/ni.3492.

18. Fletcher AL, Acton SE, Knoblich K. Lymph node fibroblastic reticular cells in health and disease. Nat Rev Immunol 2015;15(6):350–61 doi 10.1038/nri3846.

19. Acton SE, Farrugia AJ, Astarita JL, Mourao-Sa D, Jenkins RP, Nye E, et al. Dendritic cells control fibroblastic reticular network tension and lymph node expansion. Nature 2014;514(7523):498–502 doi 10.1038/nature13814.

20. Astarita JL, Cremasco V, Fu J, Darnell MC, Peck JR, Nieves-Bonilla JM, et al. The CLEC-2-podoplanin axis controls the contractility of fibroblastic reticular cells and lymph node microarchitecture. Nat Immunol 2015;16(1):75–84 doi 10.1038/ni.3035.

21. Malhotra D, Fletcher AL, Astarita J, Lukacs-Kornek V, Tayalia P, Gonzalez SF, et al. Transcriptional profiling of stroma from inflamed and resting lymph nodes defines immunological hallmarks. Nat Immunol 2012;13(5):499–510 doi 10.1038/ni.2262.

22. Martin-Villar E, Megias D, Castel S, Yurrita MM, Vilaro S, Quintanilla M. Podoplanin binds ERM proteins to activate RhoA and promote epithelial-mesenchymal transition. J Cell Sci 2006;119(Pt 21):4541–53 doi 10.1242/jcs.03218.

23. Wicki A, Lehembre F, Wick N, Hantusch B, Kerjaschki D, Christofori G. Tumor invasion in the absence of epithelial-mesenchymal transition: podoplanin-mediated remodeling of the actin cytoskeleton. Cancer Cell 2006;9(4):261–72 doi 10.1016/j.ccr.2006.03.010.

24. Diazzi S, Tartare-Deckert S, Deckert M. Bad Neighborhood: Fibrotic Stroma as a New Player in Melanoma Resistance to Targeted Therapies. Cancers (Basel) 2020;12(6) doi 10.3390/cancers12061364.

25. Kalluri R. The biology and function of fibroblasts in cancer. Nat Rev Cancer 2016;16(9):582–98 doi 10.1038/nrc.2016.73.

26. Gaggioli C, Hooper S, Hidalgo-Carcedo C, Grosse R, Marshall JF, Harrington K, et al. Fibroblast-led collective invasion of carcinoma cells with differing roles for RhoGTPases in leading and following cells. Nat Cell Biol 2007;9(12):1392–400 doi 10.1038/ncb1658.

27. Calvo F, Ege N, Grande-Garcia A, Hooper S, Jenkins RP, Chaudhry SI, et al. Mechanotransduction and YAP-dependent matrix remodelling is required for the generation and maintenance of cancer-associated fibroblasts. Nat Cell Biol 2013;15(6):637–46 doi 10.1038/ncb2756.

28. Dupont S, Morsut L, Aragona M, Enzo E, Giulitti S, Cordenonsi M, et al. Role of YAP/TAZ in mechanotransduction. Nature 2011;474(7350):179–83 doi 10.1038/nature10137.

29. Albrengues J, Bourget I, Pons C, Butet V, Hofman P, Tartare-Deckert S, et al. LIF mediates proinvasive activation of stromal fibroblasts in cancer. Cell Rep 2014;7(5):1664–78 doi 10.1016/j.celrep.2014.04.036.

30. Sanz-Moreno V, Gaggioli C, Yeo M, Albrengues J, Wallberg F, Viros A, et al. ROCK and JAK1 signaling cooperate to control actomyosin contractility in tumor cells and stroma. Cancer Cell 2011;20(2):229–45 doi 10.1016/j.ccr.2011.06.018.

31. Albrengues J, Bertero T, Grasset E, Bonan S, Maiel M, Bourget I, et al. Epigenetic switch drives the conversion of fibroblasts into proinvasive cancer-associated fibroblasts. Nat Commun 2015;6:10204 doi 10.1038/ncomms10204.

32. Robert G, Gaggioli C, Bailet O, Chavey C, Abbe P, Aberdam E, et al. SPARC represses E-cadherin and induces mesenchymal transition during melanoma development. Cancer Res 2006;66(15):7516–23 doi 10.1158/0008-5472.CAN-05-3189.

33. Didier R, Mallavialle A, Ben Jouira R, Domdom MA, Tichet M, Auberger P, et al. Targeting the Proteasome-Associated Deubiquitinating Enzyme USP14 Impairs Melanoma Cell Survival and Overcomes Resistance to MAPK-Targeting Therapies. Mol Cancer Ther 2018; 17(7): 1416–29 doi 10.1158/1535-7163.MCT-17-0919.

34. Rathore M, Girard C, Ohanna M, Tichet M, Ben Jouira R, Garcia E, et al. Cancer cell-derived long pentraxin 3 (PTX3) promotes melanoma migration through a toll-like receptor 4 (TLR4)/NF-kappaB signaling pathway. Oncogene 2019;38(30):5873–89 doi 10.1038/s41388-019-0848-9.

35. Tichet M, Prod’Homme V, Fenouille N, Ambrosetti D, Mallavialle A, Cerezo M, et al. Tumour-derived SPARC drives vascular permeability and extravasation through endothelial VCAM1 signalling to promote metastasis. Nature Communications 2015;6 doi 10.1038/ncomms7993.

36. Lino Cardenas CL, Henaoui IS, Courcot E, Roderburg C, Cauffiez C, Aubert S, et al. miR-199a-5p Is upregulated during fibrogenic response to tissue injury and mediates TGFbeta-induced lung fibroblast activation by targeting caveolin-1. PLoS Genet 2013;9(2):e1003291 doi 10.1371/journal.pgen.1003291.

37. Rodda LB, Lu E, Bennett ML, Sokol CL, Wang X, Luther SA, et al. Single-Cell RNA Sequencing of Lymph Node Stromal Cells Reveals Niche-Associated Heterogeneity. Immunity 2018;48(5):1014–28 e6 doi 10.1016/j.immuni.2018.04.006.

38. Hinz B, Celetta G, Tomasek JJ, Gabbiani G, Chaponnier C. Alpha-smooth muscle actin expression upregulates fibroblast contractile activity. Mol Biol Cell 2001;12(9):2730–41 doi 10.1091/mbc.12.9.2730.

39. Yang CY, Vogt TK, Favre S, Scarpellino L, Huang HY, Tacchini-Cottier F, et al. Trapping of naive lymphocytes triggers rapid growth and remodeling of the fibroblast network in reactive murine lymph nodes. Proc Natl Acad Sci U S A 2014; 111(1):E109–18 doi 10.1073/pnas.1312585111.

40. Ren JP, Ying RS, Cheng YQ, Wang L, El Gazzar M, Li GY, et al. HCV-induced miR146a controls SOCS1/STAT3 and cytokine expression in monocytes to promote regulatory T-cell development. J Viral Hepat 2016;23(10):755–66 doi 10.1111/jvh.12537.

41. Starr R, Willson TA, Viney EM, Murray LJ, Rayner JR, Jenkins BJ, et al. A family of cytokine-inducible inhibitors of signalling. Nature 1997;387(6636):917–21 doi 10.1038/43206.

42. Shields BJ, Hauser C, Bukczynska PE, Court NW, Tiganis T. DNA replication stalling attenuates tyrosine kinase signaling to suppress S phase progression. Cancer Cell 2008;14(2):166–79 doi 10.1016/j.ccr.2008.06.003.

43. Kim HS, Li A, Ahn S, Song H, Zhang W. Inositol Polyphosphate-5-Phosphatase F (INPP5F) inhibits STAT3 activity and suppresses gliomas tumorigenicity. Sci Rep 2014;4:7330 doi 10.1038/srep07330.

44. Kan S, Konishi E, Arita T, Ikemoto C, Takenaka H, Yanagisawa A, et al. Podoplanin expression in cancer-associated fibroblasts predicts aggressive behavior in melanoma. J Cutan Pathol 2014;41(7):561–7 doi 10.1111/cup.12322.

45. Quintanilla M, Montero-Montero L, Renart J, Martin-Villar E. Podoplanin in Inflammation and Cancer. Int J Mol Sci 2019;20(3) doi 10.3390/ijms20030707.

46. Pelon F, Bourachot B, Kieffer Y, Magagna I, Mermet-Meillon F, Bonnet I, et al. Cancer-associated fibroblast heterogeneity in axillary lymph nodes drives metastases in breast cancer through complementary mechanisms. Nat Commun 2020; 11 (1):404 doi 10.1038/s41467-019-14134-w.

47. Choi SY, Bae H, Jeong SH, Park I, Cho H, Hong SP, et al. YAP/TAZ direct commitment and maturation of lymph node fibroblastic reticular cells. Nat Commun 2020; 11(1):519 doi 10.1038/s41467-020-14293-1.

48. Haan S, Margue C, Engrand A, Rolvering C, Schmitz-Van de Leur H, Heinrich PC, et al. Dual role of the Jak1 FERM and kinase domains in cytokine receptor binding and in stimulation-dependent Jak activation. J Immunol 2008;180(2):998–1007 doi 10.4049/jimmunol.180.2.998.

49. Melnikova VO, Bar-Eli M. Inflammation and melanoma metastasis. Pigment Cell Melanoma Res 2009;22(3):257–67 doi 10.1111/j.1755-148X.2009.00570.x.

50. Berestjuk I, Lecacheur M, Diazzi S, Rovera C, Prod’homme V, Mallavialle A, et al. Targeting DDR1 and DDR2 overcomes matrix-mediated melanoma cell adaptation to BRAF-targeted therapy. bioRxiv 2019 doi 10.1101/857896.

