## Supplemental figures for "Invasive dedifferentiated melanoma cells inhibit JAK1-STAT3-driven actomyosin contractility of human fibroblastic reticular cells of the lymph node"

### Fig S1

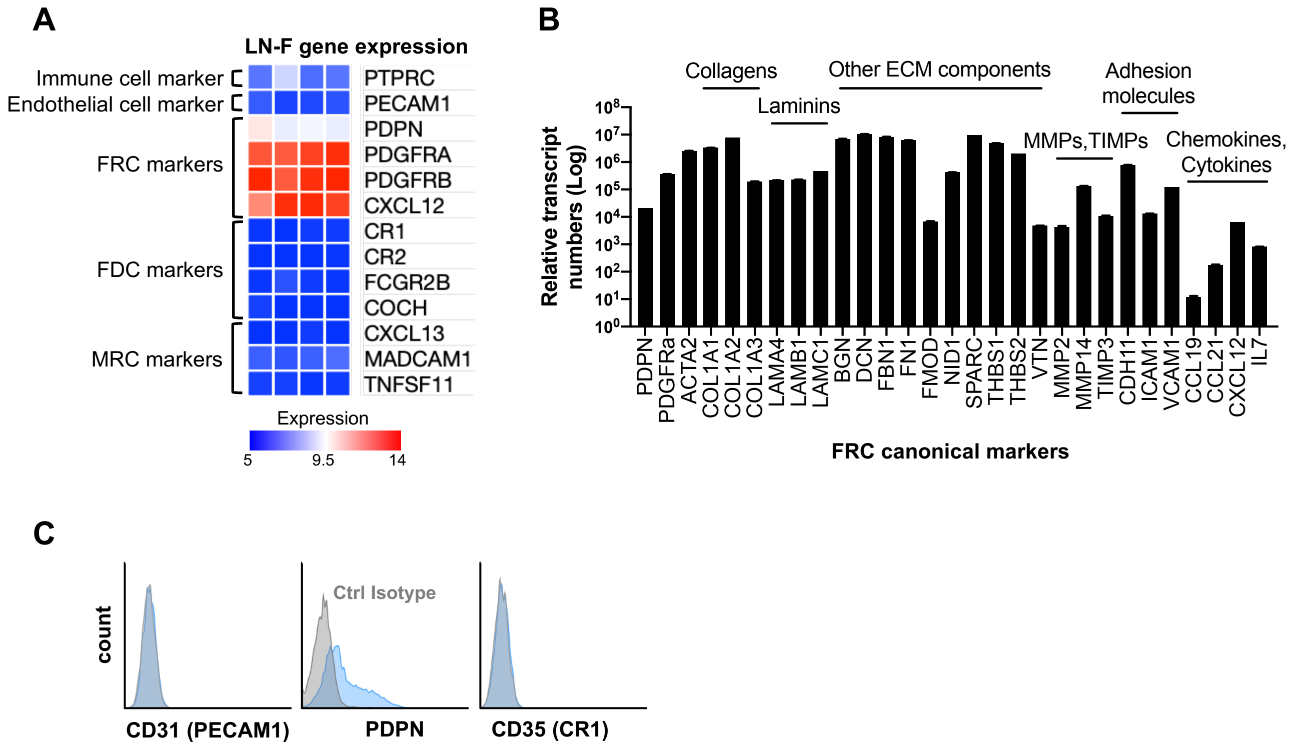


**Figure S1. LN-F express the Fibroblastic Reticular Cell (FRC) markers.**
(A) Microarray-based gene expression analysis of LN-F (quadriplicate) showing canonical markers expressed by the main LN cell subsets: immune cells, endothelial cells, FRC, Follicular Dendritic Cells (FDC) and Marginal Reticular Cells (MRC).
(B) qRT-PCR analysis of the expression of FRC-specific markers by LN-F (n = 2, in duplicate; Mean +/- SEM).
(C) Flow cytometry analysis of the expression of CD31 (PECAM1), PDPN and CD35 (CR1) on LN-F (in blue, n = 3). Staining with a control isotype mAb is shown (in grey).

### Fig S2

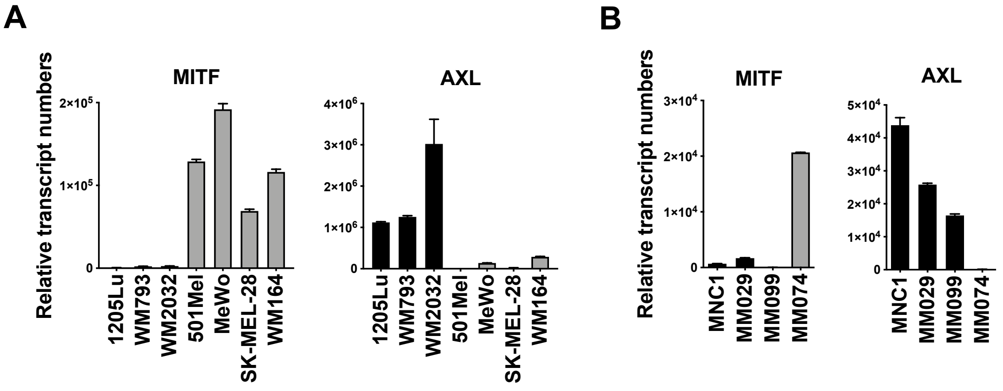


**Figure S2. Characterization of the melanocytic / dedifferentiated signature of melanoma cells.**
qRT-PCR analysis of MITF and AXL gene expression in (A) melanoma cell lines or (B) short-term melanoma cells isolated from patients (n = 2, in duplicate; mean +/- SEM).

### Fig S3

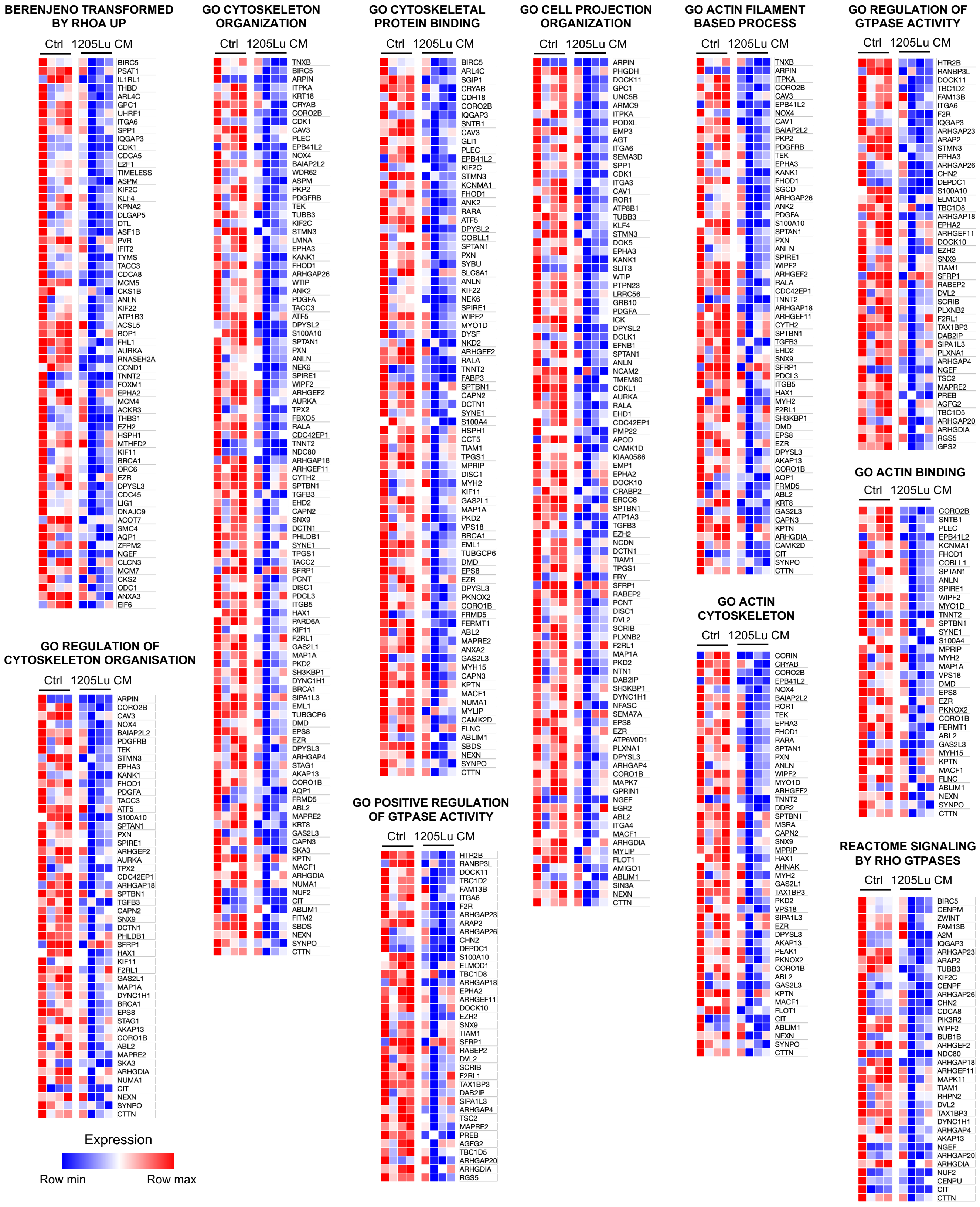


**Figure S3. Gene Set Enrichment Analysis (GSEA) of genes down-regulated in LN-F treated with 1205Lu CM compared to Ctrl LN-F.**

Genes expressed by Ctrl LN-F and LN-F treated for 48h with 1205Lu CM were compared by microarray analysis. Genes specifically down-regulated in 1205Lu CM-treated LN-F with LogFC ≤ - 0.5 were analyzed by GSEA. Pathways with the higher relative enrichment and involved in fibroblast contraction are shown.

### Fig S4

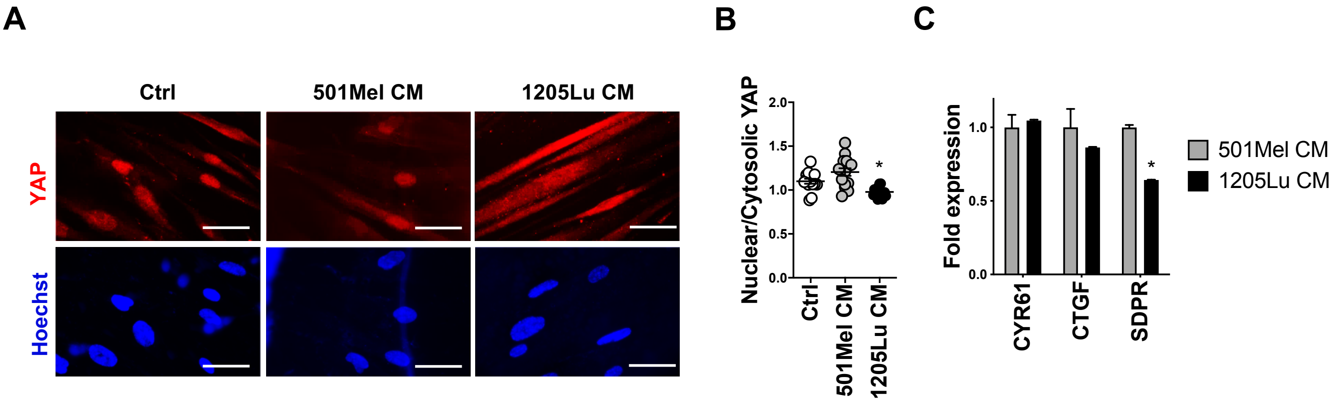


**Figure S4. 1205Lu-mediated inhibition of LN-F contraction is associated with decreased YAP activity.**
(A) Immunofluorescence analysis of YAP and Hoechst localization in LN-F plated on 0.2 kPa hydrogels and treated 4 days with 501Mel CM or 1205Lu CM (Scale bar = 50 µM).
(B) Quantification of YAP nuclear and cytosolic localization in LN-F plated on 0.2 kPa hydrogels and treated 4 days with 501Mel CM or 1205Lu CM (n = 15 cells, mean +/- SEM, p-Val (*)<0.05).
(C) Quantification by qRT-PCR of the expression of YAP target genes by LN-F plated on 0.2 kPa hydrogels and treated 4 days with with 501Mel CM or 1205Lu CM (n = 2 in duplicate, mean +/- SEM, p-Val (*)<0.05).

### Fig S5

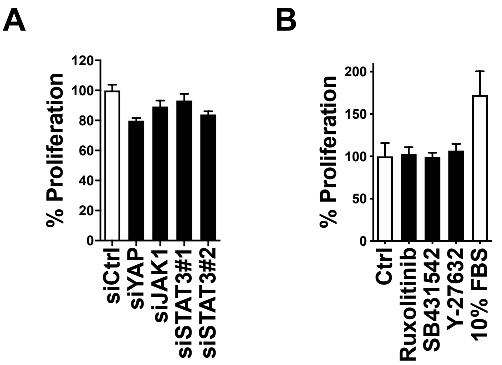


**Figure S5. Drugs and siRNAs used in gel contraction assays did not affect LN-F survival.**
(A) Proliferation for 6 days of LN-F transfected with ctrl siRNA or siRNAs depleting JAK1 (siJAK1), STAT3 (siSTAT3#1 and #2) or YAP (siYAP), expression (n = 2, in duplicate, mean +/- SEM).
(B) Proliferation of LN-F incubated 7 days with Ruxolitinib (10 µM), SB431542 (10 µM) or Y-27632 (10 µM) (n = 2 in quadruplicate, mean +/- SEM).
